# Evolution of pathogenic and nonpathogenic yeasts mitochondrial genomes inferred by supertrees and supermatrices with divergence estimates based on relaxed molecular clocks

**DOI:** 10.1101/486944

**Authors:** Luciano R. Lopes, Renata C. Ferreira, Fernando Antoneli, Paulo B. Paiva, Marcelo R. S. Briones

## Abstract

The evolution of mitochondrial genomes is essential for the adaptation of yeasts to the variation of environmental levels of oxygen. Although *Saccharomyces cerevisiae* mitochondrial DNA lacks all complex I genes, respiration is possible because alternative NADH dehydrogenases are encoded by *NDE1* and *NDI1* nuclear genes. The proposed whole genome duplication (WGD) in the yeast ancestor at 150-100 million years ago caused nuclear gene duplications and secondary losses, although its relation to the loss of complex I mitocondrial is unknown. Here we present phylogenomic supertrees and supermatrix tree of 46 mitochondrial genomes showing that the loss of complex I predates WGD and occurred independently in the *S. cerevisiae* group and the fission yeast *Schizosaccharomyces pombe*. We also show that the branching patterns do not differ dramatically in supertrees and supermatrix phylogenies. Our inferences indicated consistent relations between conserved mitochondrial chromosomal gene order (synteny) in closely related yeasts. Correlation of mitochondrial molecular clock estimates and atmospheric oxygen variation in the Phanerozoic suggests that the *Saccharomyces* lineage might have lost the complex I during hypoxic periods near Perminian-Triassic or Triassic-Jurassic mass extinction events, while the *Schizosaccharomyces* lineage possibly lost the complex I during hypoxic environment periods during Middle Cambrian until Lower Devonian. The loss of mitochondrial complex I during low oxygen might not affect yeast metabolism due to fermentative switch. The return to increased oxygen periods might have favored adaptations to aerobic metabolism. Additionally, we also showed that *NDE1* and *NDI1* phylogenies indicate evolutionary convergence in yeasts where mitochondrial complex I is absent.

## 1. Introduction

Ascomycota yeast group includes *Saccharomyces cerevisiae* and *Schizosaccharomyces pombe*, both fundamental models for eukaryotic genetics and biotechnology (Botstein et al., 1997; Hoffman et al., 2015). Despite the phylogenetic distance, *Saccharomyces* and *Schizosaccharomyces* genders share similar adaptations. In both organisms the ability to adapt to varying oxygen concentrations is essential for survival. *S. cerevisiae* and *Sz. pombe* readily ferment sugars and produce ethanol in the presence of oxygen, a process known as Crabtree effect (Crabtree, 1928; Dashko et al., 2014; Snoek and Steensma, 2006), and are also promptly able to sustained anaerobic growth despite higher energetic costs (Dashko et al., 2014; Lai et al., 2006). Therefore, these yeast species are adapted to current oxygen atmospheric levels conditions (21%) and hypoxic environments, where oxygen concentrations are lower than atmospheric levels (Postmus et al., 2011a).

During of a series of geological periods, the oxygen levels significantly varied and might be exerted a strong pressure on different species for the evolution of respiratory mechanisms and adaptations to cope with hyperoxic periods (Carboniferous, Early Perminian and Late Cretaceous-Early Tertiary), hypoxic periods (Ordovician, Devonian, Late-Perminian and Triassic-Jurassic boundary) and normoxic period (present) (Berner, 1999; Glasspool and Scott, 2010a; Ward, 2006).

The evolution of respiratory mechanisms in fungi produced patterns with different combinations of fermentation types (Li et al., 2011). For example, *Candida albicans*, the causative agent of candidiasis, the most common fungal infection worldwide, is dependent on respiration, but is able to adapt on hypoxic environment, inside its human host (Grahl et al., 2012). *C. albicans* has a complete gene set of the mitochondrial respiratory chain (Cavalheiro et al., 2004). On the other hand, *S. cerevisiae* and *Sz. pombe* lack all seven mitochondrial genes encoding for respiratory complex I subunits, but are adapted on any oxygenic conditions (Bullerwell et al., 2003; Skoneczna et al., 2015).

In *S. cerevisiae* the absence of mitochondrial respiratory complex I is compensated by *NDE1*, *NDE2* and *NDI1* nuclear gene products, although these proteins are not evolutionarily related to any of classical complex I subunits (Small and McAlister-Henn 1998; Luttik et al. 1998; Overkamp et al. 2000; Li et al. 2006). *NDE1*, *NDE2* and NDI1 proteins act transferring electrons to ubiquinone, as an alternative NADH dehydrogenase (Luttik et al., 1998; Postmus et al., 2011b). Therefore, *NDE1*, *NDE2* and NDI1 are essential for *S. cerevisiae* viability (Melo et al., 2004).

The loss of complex I genes promotes gene order variability from *Saccharomycetes* and *Schizossacharomyces* mitochondrial genomes (Aguileta et al., 2014). Fungal mitochondrial gene order is relatively free to vary, specially within Saccharomycetes (Aguileta et al., 2014). As expected, conservation of gene order in mitochondrial and nuclear genomes is higher in close related species (Keogh et al., 1998). While the loss of complex I genes is associated with the lack of conserved mitochondrial gene orders, the whole genome duplication (WGD) is associated with lack of gene orders of yeast nuclear genome (Wolfe, 2015). There is evidence that a whole genome duplication (WGD) event happened in the yeast ancestor about 150-100 million years ago (Wolfe and Shields, 1997). This genomic event doubled the dose of each nuclear gene per cell, causing gene duplications, secondary losses, neofunctionalization and the origin of ohnologs (Kellis et al., 2004; Sémon and Wolfe, 2007).

In the present study we investigated the mitochondrial phylogenomic of yeasts and whether their ability to adapt to varying oxygen concentrations could be correlated with the evolutionary history of Saccharomycetales mitochondrial genomes during the Phanerozoic (from 541 Myr to present). The great diversification of these fungi, which occurred in this geologic eon, paralleled major fluctuations in atmospheric O_2_ and CO_2_ concentrations (Berner, 1999; Glasspool and Scott, 2010a; Mentel and Martin, 2008; Ward, 2006). For this we combined phylogenomics with quantitative synteny analysis and molecular clock methods to estimate the dates of the loss of mitochondrial complex I in the *Saccharomyces* and *Schizosaccharomyces* lineages.

## 2. Materials and methods

### 2.1. Sequence Data

The thirty nine yeast complete mitochondrial genome sequences, and all other individual gene sequences, were downloaded from GenBank (http://ncbi.nlm.nih.gov/Genbank) and the Candida Genome Database (http://www.candidagenome.org), with accession numbers available in Supplementary Table 1. Individual mitochondrial gene sequences were analyzed separately. The set of mitochondrial genes including apocytochrome b (*COB*), ATP synthetase subunits (*ATP6*, *ATP8*, and *ATP9*) and cytochrome oxidase subunits (*COX1*, *COX2*, and *COX3*), conserved in genomes of all yeast species, were included in this study. Another set of (variable) genes, not present in all mitochondrial genomes, is composed by NADH dehydrogenase (ubiquinone) complex I subunits (*NAD1*, *NAD2*, *NAD3*, *NAD4*, *NAD4L*, *NAD5* and *NAD6*). Additionally, large (LSU) and small (SSU) ribosomal RNA subunits genes were also analyzed. Sequences were aligned, individually, using Clustal W (Larkin et al., 2007), with default settings, and manually adjusted using Seaview software (Galtier et al., 1996).

**Table 1.**
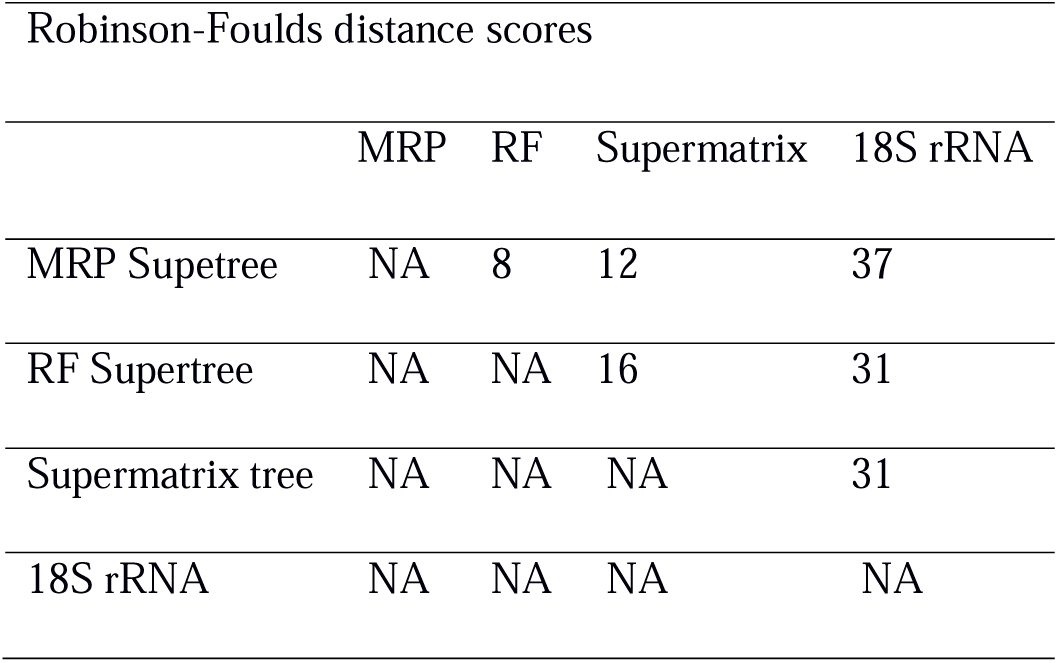
Robinson Foulds distance scores of supertrees.

|  | MRP | RF | Supermatrix | 18S rRNA |
| --- | --- | --- | --- | --- |
| MRP Supetree | NA | 8 | 12 | 37 |
| RF Supertree | NA | NA | 16 | 31 |
| Supermatrix tree | NA | NA | NA | 31 |
| 18S rRNA | NA | NA | NA | NA |

### 2.2. Inference of Source Trees

To infer phylogenetic Bayesian trees, substitution models were determined for each dataset based on the Akaike Information Criterion (AIC) and Bayesian Information Criterion (BIC), using MEGA 6 software (“find best DNA model” option) (Tamura et al., 2013). Depending on the gene, we selected either general time-reversible (GTR) model (Rodríguez et al., 1990) with Γ distribution to correct for rate variation among sites (GTR+G) or GTR with Γ distribution and proportion of invariant sites, (GTR+G+I) (Yang, 1994) (the complete parameter list is available in Supplementary Table 2).

**Table 2.**
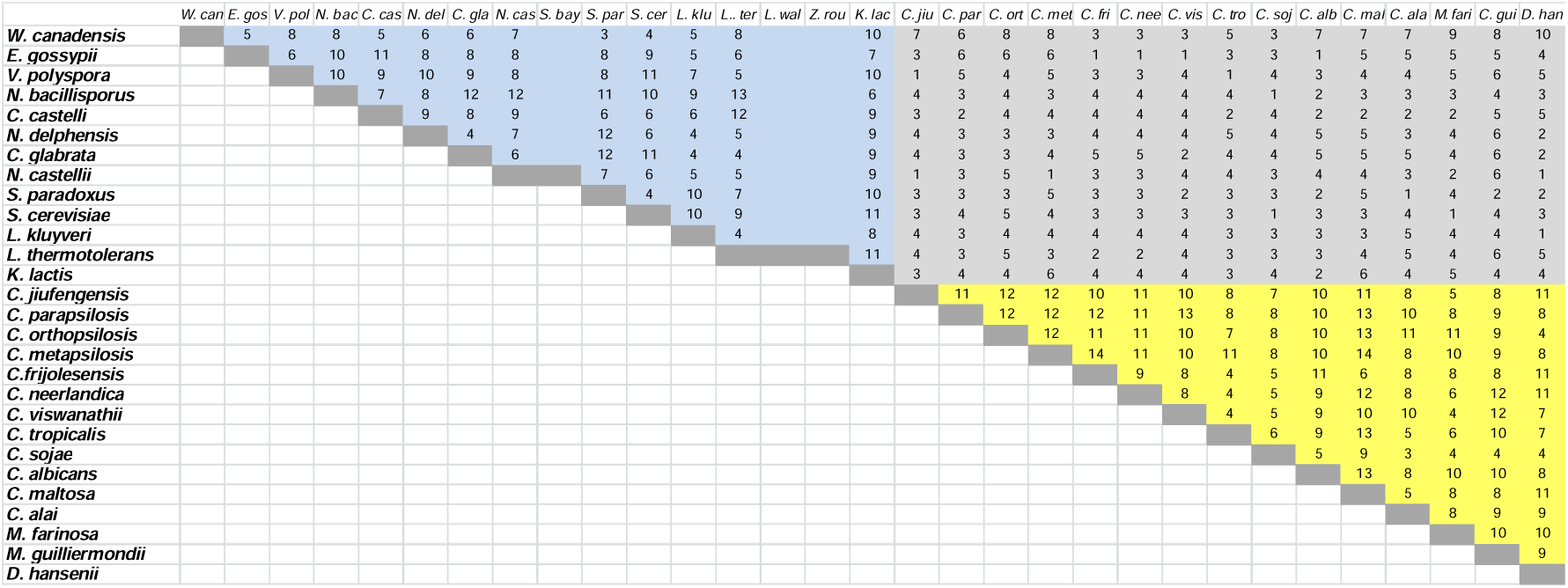
Quantitative synteny of yeast mitochondrial genomes.

All the mitochondrial source trees and the nuclear phylogenies based on 18S rRNA, *NDE1* and *NDI1* genes were inferred using MrBayes 3.2 software (Ronquist et al., 2012). The gene sequences were downloaded from GenBank according to the availability. Only 37 nucleotide sequences of 18S rRNA gene from the species set here considered were available, while few species had *NDE1* and NDI1 sequences available (see Supplementary Table 1). Bayesian trees were searched for 10^7^ generations with sampling every 100 generations until the standard deviation from split frequencies were under 0.01. The parameters and the trees were summarized by wasting 25% of the samples obtained (burn-in). The consensus trees were then used to determine the posterior probabilities. All phylogenetic trees were then formatted with the FigTree v1.3.1 software (http://tree.bio.ed.ac.uk/software/figtree/).

### 2.3. Inference of Supertree and Supermatrix

The 16 mitochondrial source trees (Supplementary Figures 1-16) were combined as a comprehensive Robinson-Foulds (RF) supertree and also as a Matrix Representation Parsimony (MRP) supertree. The mitochondrial RF supertree was inferred using RF-supertree software (http://genome.cs.iastate.edu/CBL/RFsupertrees) and CompPHY software (http://www.atgc-montpellier.fr/compphy/) was used to infer MRP supertree (Fiorini et al., 2014). Both supertree methods consider even the phylogenetic source trees with low frequency species overlap to generate a supertree containing all species. The analysis included genomes with varying gene contents and three species (*Lachancea waltii*, *Saccharomyces bayanus* and *Zygosaccharomyces rouxii*) represented only by 15S rRNA and COX2 gene, because their mitochondrial genomes were not available.

**Figure 1.**
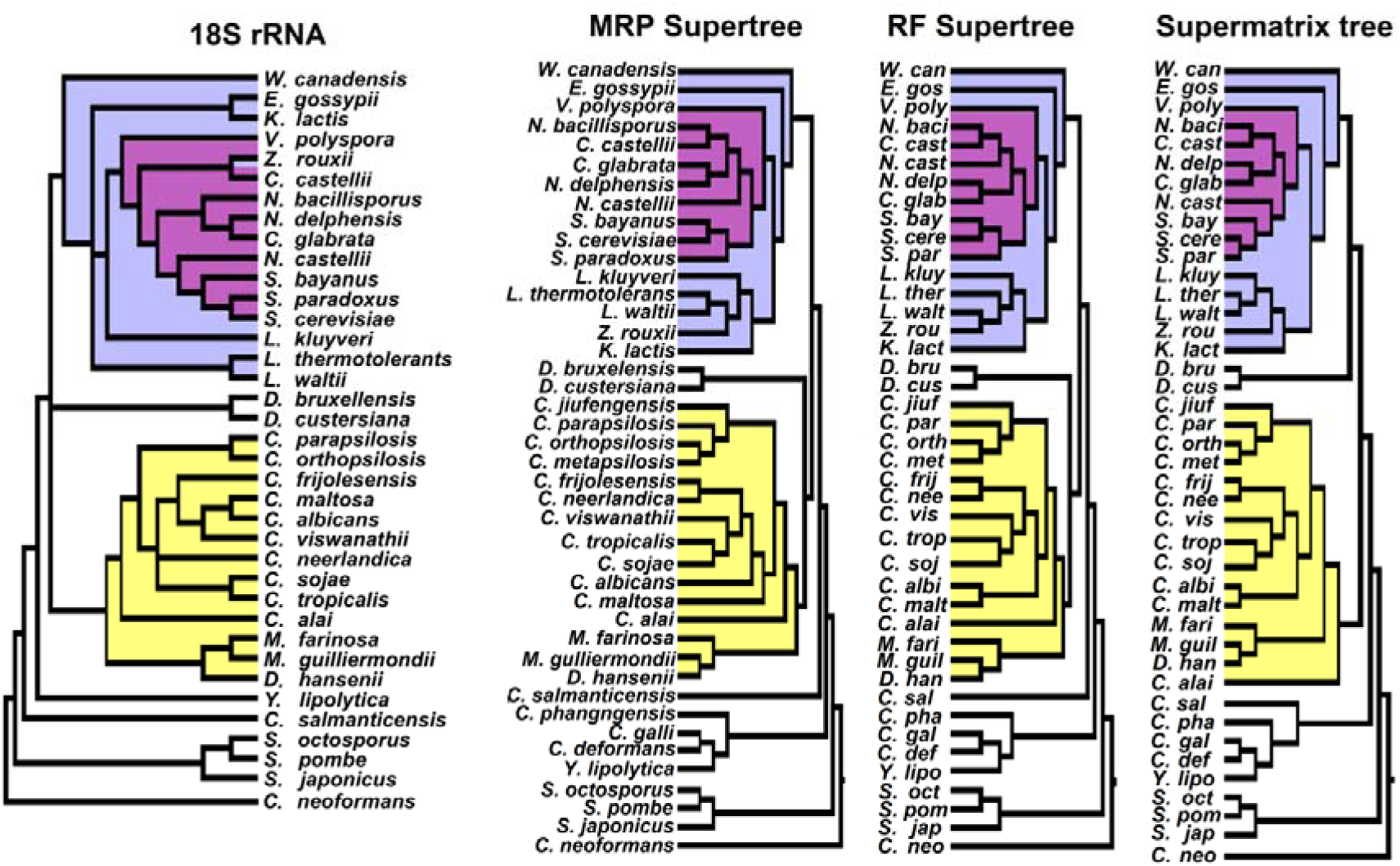
Multigene phylogenetic analysis, supermatrix phylogenetic tree and supertrees. Supertrees were inferred from 16 source trees, each corresponding to all Saccharomycota mitochondrial genes excepting the tRNA genes. Phylogenetic tree based on supermatrix was constructed using concatenated alignment of mitochondrial gene sequences. Only three node labels had posterior probabilities values inferior than 1. RF Supertree was inferred using the Robinson-Foulds algorithm as implemented in the RF program (Bansal et al., 2010). Supertree based on Matrix Representation Parsimony algorithm (Baum, 1992; Ragan, 1992), MRP Supertree, was constructed using CompPHY software (Fiorini et al., 2014). Phylogenetic tree based on 18S rRNA was the only tree inferred using single gene sequence. The blue group includes pre and pos-WGD species. WGD was highlighted in purple. The yellow group includes most Candida species, the CTG clade.

The RF supertree provide a binary supertree, by heuristic search, that minimizes the sum of the Robinson-Foulds distances (defined as a normalized count of the symmetric difference between the sets of clusters of the two trees) between every rooted source tree and the supertree (Bansal et al. 2010). Fourty four optimal trees output were generated by RF-supertree software with default set. All the optima supertrees were combined into a single consensus tree using Geneious v6.1.6 software.

MRP has been the method of choice for most of the supertree analysis. The MRP method, independently developed by Baum (1992) and Ragan (1992) (Baum, 1992; Ragan, 1992), consists in constructing a matrix where each taxon is coded according to the presence in a clade (“1” for presence, “0” for absence in the clade and “?” when absent in the source tree). Based on the constructed matrix, the most parsimonious tree was built. CompPHY software provided MRP supertree that was edited in Figtree v1.3.1.

The software based on RF and MRP are able to estimate supertrees that are consistent with the largest number of clusters (or clades) from the source trees. The approach is able to consider even the phylogenetic source trees with low frequency species overlap to generate a supertree containing all species. Species included in this study, such as *Sz. pombe*, for example, that have low number of genes in its mitochondrial genome, had not been included in all sources trees, but was included in supertrees. Even those species which were represented only by 15S rRNA and COX2 gene were included in supertrees.

The concatenated align sequences to construct the supermatrix phylogenetic tree included sixteen mitochondrial genes dataset (22.6 kilobases). Alignments, from mitochondrial genes, were concatenated using Seaview software. Best-fit models of molecular evolution for concatenated aligment were selected based on AIC and BIC using MEGA 6 software (Lanfear et al., 2012) (Supplementary Table 2). A Bayesian tree with partitioning schemes was inferred based on concatenated alignment using MrBayes 3.2 software with 10^7^ generations, sampling every 100 generations, until the split frequencies standard deviations were under 0.01. The selected trees were summarized by wasting 25% of the samples obtained (burn-in) and a consensus trees were then used to determine the posterior probabilities values.

### 2.4. Robinson-Foulds distances

To infer congruency between two phylogenetic trees, we use RF distance. The symmetric difference between the sets of clusters of the two phylogenetic trees can be defined as a normalized count by RF distance, as score value. The most congruent tree comparison have low RF-distance, low score. Using CompPHY software (Fiorini et al., 2014) to provide RF distance scores, we compare the congruency between supertrees, supermatrix phylogeny and nuclear 18S rRNA.

### 2.5. Mitochondrial Genome Synteny Analysis

The synteny (or gene order) map was constructed using Geneious v6.1.6 to verify gene positions and transcriptional directions. The graphic with gene order and gene presence/absence was constructed using Microsoft Excel and Libre Office Draw. All yeast mitochondrial genomes were delineated with all genes linearly depicted as arrow boxes indicating forward (right) or reverse (left) sense. Genomes were manually aligned, using 15S rRNA as starting point, and progressively assembly blocks of genes that exhibit conserved features across genome. Mitochondrial genomes were pairwise compared using the progressive Mauve software (Darling et al., 2010) to search for regions of local synteny (colinear blocks), counted as units of conserved blocks. The nonparametric Mann-Whitney test was used to infer that the synteny mean values are distinct among the clades, using GraphPad Instat software.

NADH gene remnants were searched, using BLAST (Altschul et al., 1990), within nuclear genome of yeasts *S. cerevisiae*, *Sz. pombe* and *C. glabrata*. The search was optimized using megablast option, to find highly similar sequences, and blastn option, for sequences with somewhat similarity.

### 2.6. Molecular Clock Tests and Divergence Date Estimates

For divergence date estimates we used BEAST v2.3.1 software (Drummond et al., 2012). The null hypothesis of a strict global clock was rejected for all mitochondrial genes, by the likelihood ratio test for α=0.05 (Felsenstein, 1981). Likelihood ratio test for molecular clock was calculated based on 2Δ, the difference between the likelihood scores (LnL□-LnL□) of the trees, considering that LnL□ represents likelihood from molecular clock tree and LnL□ represents likelihood from non-clock tree. Molecular clock only allows null hypothesis model. Non-molecular clock allows alterative hypothesis. In testing molecular clock hypothesis, the degrees of freedom work out to be s-2, where “s” is the number of taxa in the phylogeny (Felsenstein, 1981). Substitution models were selected based on AIC and BIC using MEGA 6 software for each mitochondrial gene. We implemented Yule speciation process prior. Bayesian coestimation of topology and divergence time were obtained using uniform prior for calibration ages. The divergence dates for the mrca priors of Ascomycota (811.57Myr and 690.55Myr) and Saccharomycetales (473.33Myr and 648.32Myr), obtained from Padovan et al. (2005) study, were used as calibration points. BEAST analyses were run for 10 million generations, logging parameters and trees every 1000 generations.

### 3. Results and Discussion

After the alignment of 39 mitochondrial complete yeast genomes (Supplementary Table 1) the coding domains sequences of 16 genes were extracted and used to infer unrooted Bayesian phylogenies for each separate gene. These trees (or source trees), available as supplementary figures 1–16, were used as inputs for the inferences of the Robinson-Foulds (RF) supertree (Bansal et al., 2010) and Matrix Representation Parsimony (MRP) supertree (Baum, 1992; Ragan, 1992), in figure 1 (and Supplementary Figures 17 and 18). RF supertree is the tree that minimizes the RF distances between each individual source tree and considers the presence-absence of species in different source trees. This supertree is estimated by the heuristic tree-bissection reconnection where the RF distances are the number of operations necessary to convert one topology to another (Bansal et al., 2010). On the other hand, MRP supertree is based on the constructed matrix where each taxon is coded according to the presence in a clade (“1”), absence in the clade (“0”) or absent in the source trees (“?”). Thus, the most parsimonious tree built represents a supertree (Bininda-Emonds, 2004; Sanderson et al., 1998).

Supertrees methods have been broadly applied in phylogenetic studies, aimed the combination of phylogenetic trees based on different types of data, joining molecular and morphology (Beck et al., 2006; Bininda-Emonds et al., 2007; Chang et al., 2013; Creevey and McInerney, 2005; Davis and Page, 2014; Duda and Jan Zrzavý, 2016; Liu et al., 2001; Nyakatura and Bininda-Emonds, 2012; Ruta et al., 2007; Sigwart and Lindberg, 2015), or exclusively molecular data (Campbell and Lapointe, 2010; Daubin et al., 2001; Fitzpatrick et al., 2006; Higdon et al., 2007; Modica et al., 2011; Swenson et al., 2011). More specifically, mitochondrial genes datasets were used to construct supertrees (Campbell and Lapointe, 2010; Fitzpatrick et al., 2006; Higdon et al., 2007).

In figure 1, we observe that MRP and RF supertrees included two groups which correspond to the four nuclear (Nc) and mitochondrial (Mt) genetic code patterns observed: Group 1 (blue group) has Nc=1 (standard) (Abramczyk et al., 2003) and Mt=3 (yeast mitochondrial) (Clark-Walker and Weiller, 1994) and Group 2 (yellow group) has Nc=1 and 12 (alternative yeast nuclear genetic code) (Ohama et al., 1993) and Mt=4. All species from blue group are included in Saccharomycetacea family plus *Wickerhamomyces. canadensis* that belongs to Wickerhamomycetaceae family. This group includes also the species that were described as WGD positive species (in purple). Both supertree methods supported similar topology, but there are few differences in blue group – in phylogenetic position of pre-WGD and post-WGD species.

Supertree methodology has been considered a robust approach to analyze broad phylogenies (Daubin et al., 2001; Qian and Zhang, 2016), but the difficulty to carry statistical analysis is considered challenging in supertree methodology (von Haeseler, 2012). Therefore, the phylogenetic approach based on Bayesian trees could compensate this disadvantage in supertree methodology. Bayesian inference allows analysis of large datasets and incorporates full models of sequence evolution and is not restricted to a unique best tree (Padovan et al., 2005). Moreover, we tried to compensate the lack of statistical support by inferring the Bayesian source trees with the best gene substitution models (Supplementary table 2), including high number of tree generations and considering significant statistical support measures to combine source trees into RF and MRP supertrees. Each source tree, obtained by MrBayes, reached estimated sample size (ESS) values above 200 (which should be at least 100), and reached the potential scale reduction factor (PSRF) values equal or above to 1.0 (recommendation is close to 1.0). Furthermore, 87.5% of posterior probabilities values of source trees were equal or higher than 0.75 (Supplementary Figures 1-16). Posterior probability represents the probability that a clade is true given the data, the model, and the priors (Larget and Simon, 1999).

In addition to supertrees, we made a supermatrix phylogenetic tree (Fig. 1). We applied yeasts mitochondrial genome sequences dataset, the same dataset used in supertrees, to infer a phylogenomic tree based on concatenated alignment (also called supermatrix). Differently from supertrees methodologies, supermatrix phylogenetic tree was not constructed by source trees combination. Instead, phylogenetic tree was constructed based on concatenated alignments (head to tail linkage) from 16 mitochondrial genes sequences. By supermatrix phylogenetic methodology, we constructed a Bayesian phylogenetic tree that allowed us to infer satisfactory statistical support values (ESS and PSRF). Mitochondrial supermatrix phylogenetic tree also presented consistent posterior probability values. Only three nodes had support values inferior to 1 (see Supplementary Figure 19).

Supermatrix phylogenetic tree also conserved clades presented in RF and MRP supertrees. Differently from supertrees, supermatrix phylogenetic tree placed *Millerozyma farinosa*, *Meyerozyma guilliermondii* and *Debaryomyces hansenii* near the other CTG clade species than *C. alai*, similarly as previous study (Valach et al., 2011). *Dekkera* gender is the first branch out from blue group, while into supertrees *Dekkera* gender is the first branch out from yellow group.

Despite some minor topologies divergences between supertrees and supermatrix tree, the pattern of branching trees did not deviate dramatically (Figure 1). Most of clades were conserved even in supertrees such as in supermatrix. As well as Nyakatura and Bininda-Emonds study (2012), our results indicated that there were not significant differences between supertrees and supermatrix topologies (Nyakatura and Bininda-Emonds, 2012), even considered higher accuracy from supermatrix method comparing with supertree (Kupczok et al., 2010).

Besides multi-gene mitochondrial phylogeny, we also included 18S rRNA phylogenetic tree (Figure 1 and Supplementary Figure 20), representing nuclear evolution hypothesis. Mitochondrial genome evolves considerably faster than nuclear genome due strong selective pressure under mitochondria (Brown et al., 1979; Saccone et al., 2000). Consequently, mitochondrial genomes from different yeast species present variable size and structure and accumulate higher mutation frequencies than nuclear genomes (Saccone et al., 2000). Nuclear 18S rRNA phylogeny has the same three groups identified in the mitochondrial phylogenies, but with additional differences along the tree (inconclusive place of *Dekkera*, inclusion of *Z. rouxii* into WGD group and polytomy into CTG clade, for example).

To compare differences into topologies we applied Robinson-Foulds distance method. This method quantifies topological differences between two phylogenetic trees. Comparison between more congruent trees results in low RF distance. Based on RF distance score, supertrees (MRP and RF) phylogeny are the more congruent trees (scores = 8, see Table 1). When nuclear 18S rRNA based tree is compared with supertrees or supermatrix the scores of RF distances are higher than comparisons between mitochondrial trees.

To ratify the phylogenetic results and support the topology of conserved groups placed on the mitochondrial phylogenetic trees, we investigated the gene order of mitochondrial genomes from yeast species comparing with phylogeny (Figure 2). MRP supertree (most congruent with other trees - see Table 1) was compared with mitochondrial gene order map which revealed that the phylogeny correspond to synteny patterns. Fungal mitochondrial genomes have highly variable size between their species, with mtDNA ranging from 19kbp in *Sz. pombe* to 100 kbp *S. cerevisiae*. Therefore, co-localization of the block of the genes into genome was quantified, using Mauve software, and is very similar between closely related yeast species (Table 2). The pairwise matrix comparison confirmed higher synteny between species within the same clade than synteny between species from different clades, supported by nonparametric Mann-Whitney (Table 3).

**Table 3.** Synteny statistics.

| Synteny means (standard deviation) |  | Diference means | Mann-Whitney test |
| --- | --- | --- | --- |
| Clade 1 | Clade 1 and 2 |  | p value |
| 7.88 (2.38) | 3.75 (1.53) | 4.13 | >0.0001 |
| Clade 2 | Clade 2 and 3 |  |  |
| 8.77 (2.63) | 3.75 (1.53) | 5.02 | >0.0001 |

**Figure 2.**
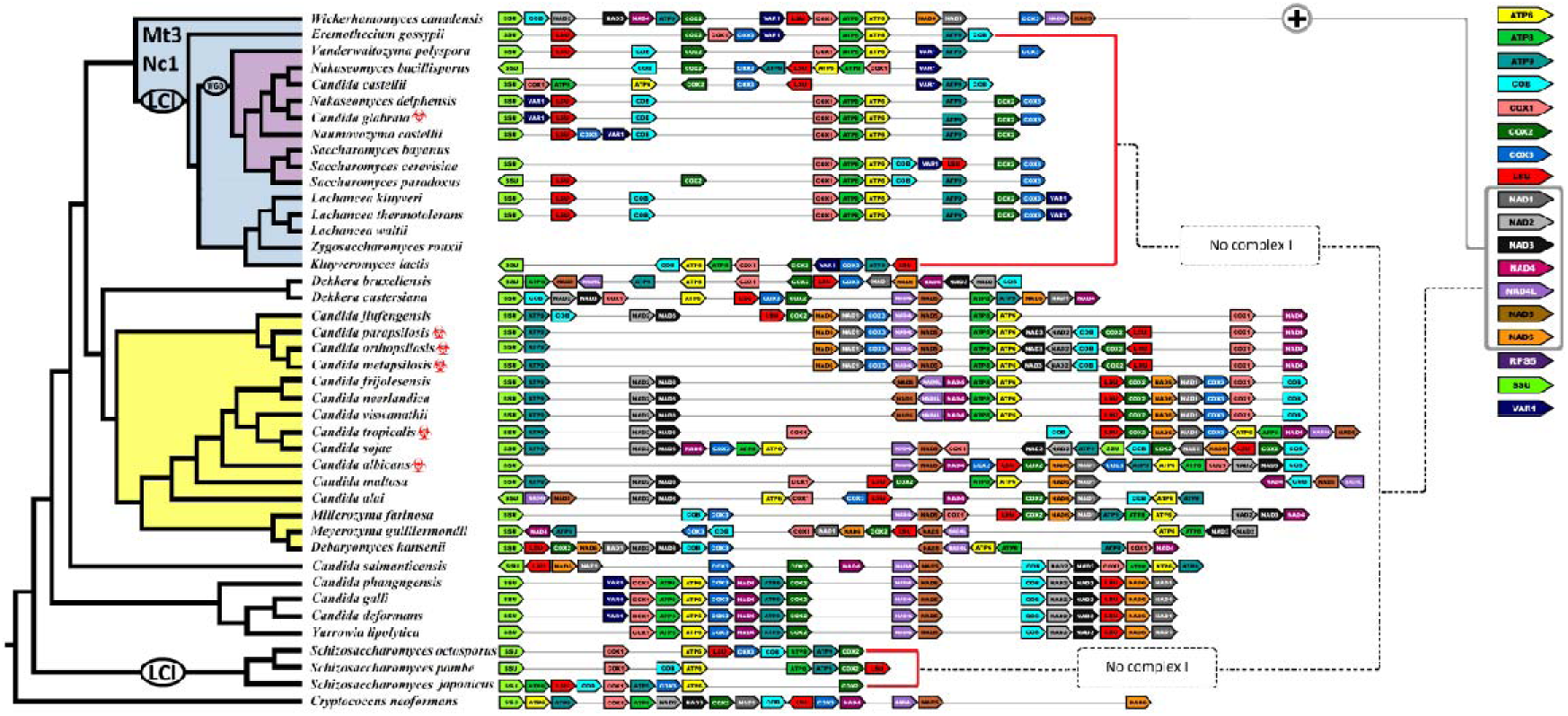
MRP supertree, most congruent tree, represents mitochondrial phylogeny. Mitochondrial gene order map of every yeast specie is represented by colored boxes with direction according to the coding sense. The blue group, (Saccharomyces lineage), has nuclear code 1 and mitochondrial code 3. WGD occurred in ancestors from the blue group, highlighted as purple clade. The gene order map indicates that the blue group has no genes encoding mitochondrial complex I (NADH dehydrogenase subunits), except for *W. canadensis*. This is an evidence that the loss of complex I occurred after the divergence between *W. canadensis* and *Saccharomyces* lineage. The yellow group contains only representatives of genus Candida, including C. albicans, with nuclear code 1 and 12 and mitochondrial 4. Biohazard symbols indicate pathogenic yeasts.

Comparison between gene order map and phylogeny allowed us to identified ancestral point from the mitochondrial complex I loss (Figure 2). This comparison also showed that the absence of mitochondrial complex I occurred independently in *Saccharomyces* lineage and the *Schizosaccharomyces* genus, also presented by previous studies (Bullerwell et al., 2003; de Zamaroczy and Bernardi, 1986; Nosek and Fukuhara, 1994; Procházka et al., 2010). After the divergence between *W. canadensis* and the common ancestral of yeast species that composed blue group (see Fig. 2) occurred an event (or a sequence of events) that promoted the loss of complex I. The loss of mitochondrial complex I in the *Saccharomyces* lineage leads us to suggest that complex I might, at least in part, affected genome stability. Nuclear genome instability may be resulted from a failure of various mitochondrial functions, such as an electron transport chain activity breakdown, for example, which changed ATP dynamic production (Kaniak-Golik and Skoneczna, 2015). Functional mitochondria are crucial to maintaining the stability of cellular genetic information (Skoneczna et al., 2015). This requirement is not limited to the proteins and RNAs encoded by the mitochondrial genome or proteins whose expression needs tight feedback control. Mitochondrial dysfunction can results in increased reactive oxygen species (ROS) production that, consequently, increases frequencies of mutations and lesions in the mitochondrial genome and may lead the loss of parts of mitochondrial DNA or all mitochondrial genome (Rasmussen et al., 2003; Yazgan and Krebs, 2012). So, mutations and deletions may have affected the mitochondrial genome and, consequently, impaired the stability of the nuclear genome (Veatch et al., 2009).

The absences of mitochondrial complex I from *S. cerevisiae* and *Sz. pombe* were documented by previous studies (Bullerwell et al., 2003; de Zamaroczy and Bernardi, 1986; Lang and Wolf, 1984; Nosek and Fukuhara, 1994; Procházka et al., 2010). However, when the loss of complex I happened remained unkown. Moreover, whether *Schizosaccharomyces* and *Saccharomyces* lineages loose the complex I in the same episode (or not) was not explored yet. For while, estimated divergence of *S. cerevisiae* and *Sz. pombe* is widely ranged between 200 Myr until 1000 Myr, and the divergence between *Candida albicans* and *S. cerevisiae* is estimated to be 871 Myr, by multiprotein analysis and 526.95 Myr depending on the calibration of 18S rRNA phylogeny (Heckman et al., 2001; Padovan et al., 2005). The mitochondrial supertrees and phylogenetic tree based on supermatrix indicate that the loss of mitochondrial complex I occurred after the divergence of *W. canadensis* from the *Saccharomyces* lineage, certainly older than 150-100 Myr, the estimate for the WGD (Wolfe and Shields, 1997). To estimate the date of complex I loss and WGD we used molecular clock analysis based on mitochondrial genes COB, ATP synthetase subunits (ATP6, ATP8, and ATP9), cytochrome oxidase subunits (COX1, COX2 e COX3) and LSU and SSU ribosomal RNA genes. We also included nuclear 18S rRNA to estimate loss of mitochondrial complex I.

Based in our dataset genes, the likelihood ratio test for molecular clock rejected the null hypothesis of a strict global clock by comparison of the likelihoods of the clock constrained tree versus the non-clock unconstrained tree (see Supplementary Table 3 for details). This approach is considered more realistic and focused on estimate periods with variable rates (Wilke et al., 2009).

Defined the model of molecular clock, we aimed to estimate when the loss of mitochondrial complex I occurred. As indicated in the phylogenetic analysis (Figure 2), the loss of mitochondrial complex I occurred after the divergence between *W. canadensis* with other species from blue group. Additionally, we also estimated loss of complex I from *Schizosaccharomyces* genus. Concomitantly, WGD, previously dated between 150 Myr to 100 Myr (Wolfe and Shields, 1997), was reanalyzed with our dataset.

For molecular clock analysis, we adopted the confidence intervals (95% HPDI -highest posterior density interval) as main result, taking into account the low reliability of mean ages. Confidence intervals may provide more likelihood dates, even when intervals are broad (Warnock et al., 2017).

In agreement with phylogenetic and synteny results, molecular clock pointed the loss of mitochondrial complex I by *Schizosaccharomyces* and *Saccharomyces* occurred in distinct events (Table 4 and Figure 4). Considering the confidence intervals based on nine mitochondrial genes, most of them (seven genes – *ATP6*, *COB*, *COX1-* 3, SSU and LSU rRNA, see Table 4 and Figure 4A), dated the loss of complex I by *Schizosaccharomyces* in a period which comprehends the middle of Cambrian until the middle of Carboniferous (Figure 4B). The estimates for the loss of complex I by *Schizosaccharomyces* based on *COB* (479.850 - 402.595 Myr), *COX3* (496.991 - 419.014 Myr), LSU (507.809 - 420.295 Myr) and SSU (459.401 - 403.040 Myr) encompass the Late Ordovician mass extinction event Figure 4B). This event, dated between 458 and 443 Myr, probably occurred due variation of climatic conditions, characterized by a high CO_2_ concentration, hypoxia and glaciation (Berner, 1999; Finnegan et al., 2011; Pohl et al., 2014; Raup and Sepkoski, 1982; Sheehan, 2001). The Late Ordovician extinction event eliminated around 25% of the families and nearly 60% of the genera of marine organisms (Wake and Vredenburg, 2008). This scenario seems to provide conditions for the loss of a mitochondrial complex I due low atmospheric oxygen availability. Alternatively, the Late Devonian mass extinction event (around 364 Myr), was encompassed in the estimate for loss of complex I by *Schizosaccharomyces* lineage, based on *ATP6* (409.057 - 330.160 Myr), *COX1* (381.190 - 314.141 Myr) and *COX2* (424.094 - 335.067 Myr). Late Devonian mass extinction event might have offered significant pressure on microbial community, as discussed above, due hypoxic ocean and atmospheric environments (Berner, 2006; McGhee, 2012). This event, which a bolide impact as a cause controversial (Racki, 2012), shifted the temperature down and decreased oxygen levels, causing major mass extinction of marine life, around 57% of marine genera (Wake and Vredenburg, 2008; Ward, 2006).

**Table 4.**
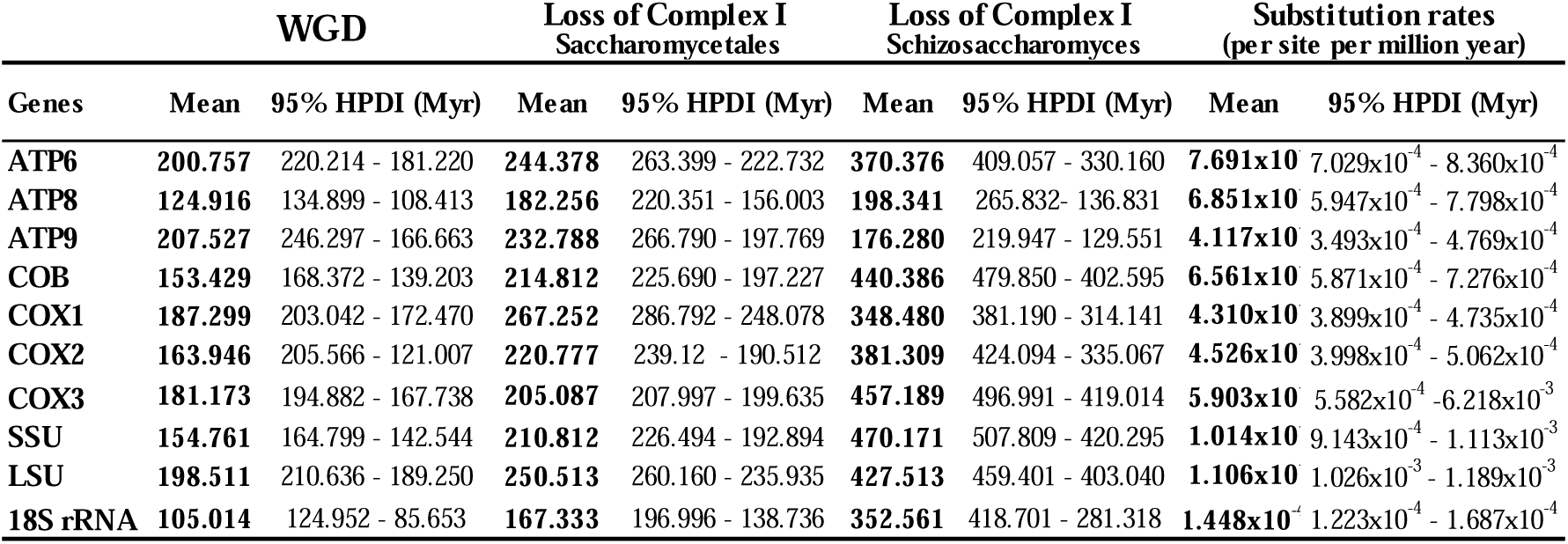
Divergence date estimates with 95% highest posterior density interval (HPDI). WGD = Whole Genome Duplication and Myr = Million years.

| Genes | WGD |  | Loss of Complex I<br>Saccharomycetales |  | Loss of Complex I<br>Schizosaccharomyces |  | Substitution rates<br>(per site per million year) |  |
| --- | --- | --- | --- | --- | --- | --- | --- | --- |
|  | Mean | 95% HPDI (Myr) | Mean | 95% HPDI (Myr) | Mean | 95% HPDI (Myr) | Mean | 95% HPDI (Myr) |
| ATP6 | **200.757** | 220.214 - 181.220 | **244.378** | 263.399 - 222.732 | **370.376** | 409.057 - 330.160 | **7.691x10<sup>-4</sup>** | 7.029x10<sup>-4</sup> - 8.360x10<sup>-4</sup> |
| ATP8 | **124.916** | 134.899 - 108.413 | **182.256** | 220.351 - 156.003 | **198.341** | 265.832 - 136.831 | **6.851x10<sup>-4</sup>** | 5.947x10<sup>-4</sup> - 7.798x10<sup>-4</sup> |
| ATP9 | **207.527** | 246.297 - 166.663 | **232.788** | 266.790 - 197.769 | **176.280** | 219.947 - 129.551 | **4.117x10<sup>-4</sup>** | 3.493x10<sup>-4</sup> - 4.769x10<sup>-4</sup> |
| COB | **153.429** | 168.372 - 139.203 | **214.812** | 225.690 - 197.227 | **440.386** | 479.850 - 402.595 | **6.561x10<sup>-4</sup>** | 5.871x10<sup>-4</sup> - 7.276x10<sup>-4</sup> |
| COX1 | **187.299** | 203.042 - 172.470 | **267.252** | 286.792 - 248.078 | **348.480** | 381.190 - 314.141 | **4.310x10<sup>-4</sup>** | 3.899x10<sup>-4</sup> - 4.735x10<sup>-4</sup> |
| COX2 | **163.946** | 205.566 - 121.007 | **220.777** | 239.12 - 190.512 | **381.309** | 424.094 - 335.067 | **4.526x10<sup>-4</sup>** | 3.998x10<sup>-4</sup> - 5.062x10<sup>-4</sup> |
| COX3 | **181.173** | 194.882 - 167.738 | **205.087** | 207.997 - 199.635 | **457.189** | 496.991 - 419.014 | **5.903x10<sup>-4</sup>** | 5.582x10<sup>-4</sup> - 6.218x10<sup>-3</sup> |
| SSU | **154.761** | 164.799 - 142.544 | **210.812** | 226.494 - 192.894 | **470.171** | 507.809 - 420.295 | **1.014x10<sup>-3</sup>** | 9.143x10<sup>-4</sup> - 1.113x10<sup>-3</sup> |
| LSU | **198.511** | 210.636 - 189.250 | **250.513** | 260.160 - 235.935 | **427.513** | 459.401 - 403.040 | **1.106x10<sup>-3</sup>** | 1.026x10<sup>-3</sup> - 1.189x10<sup>-3</sup> |
| **18S rRNA** | **105.014** | 124.952 - 85.653 | **167.333** | 196.996 - 138.736 | **352.561** | 418.701 - 281.318 | **1.448x10<sup>-4</sup>** | 1.223x10<sup>-4</sup> - 1.687x10<sup>-4</sup> |

**Figure 3.**
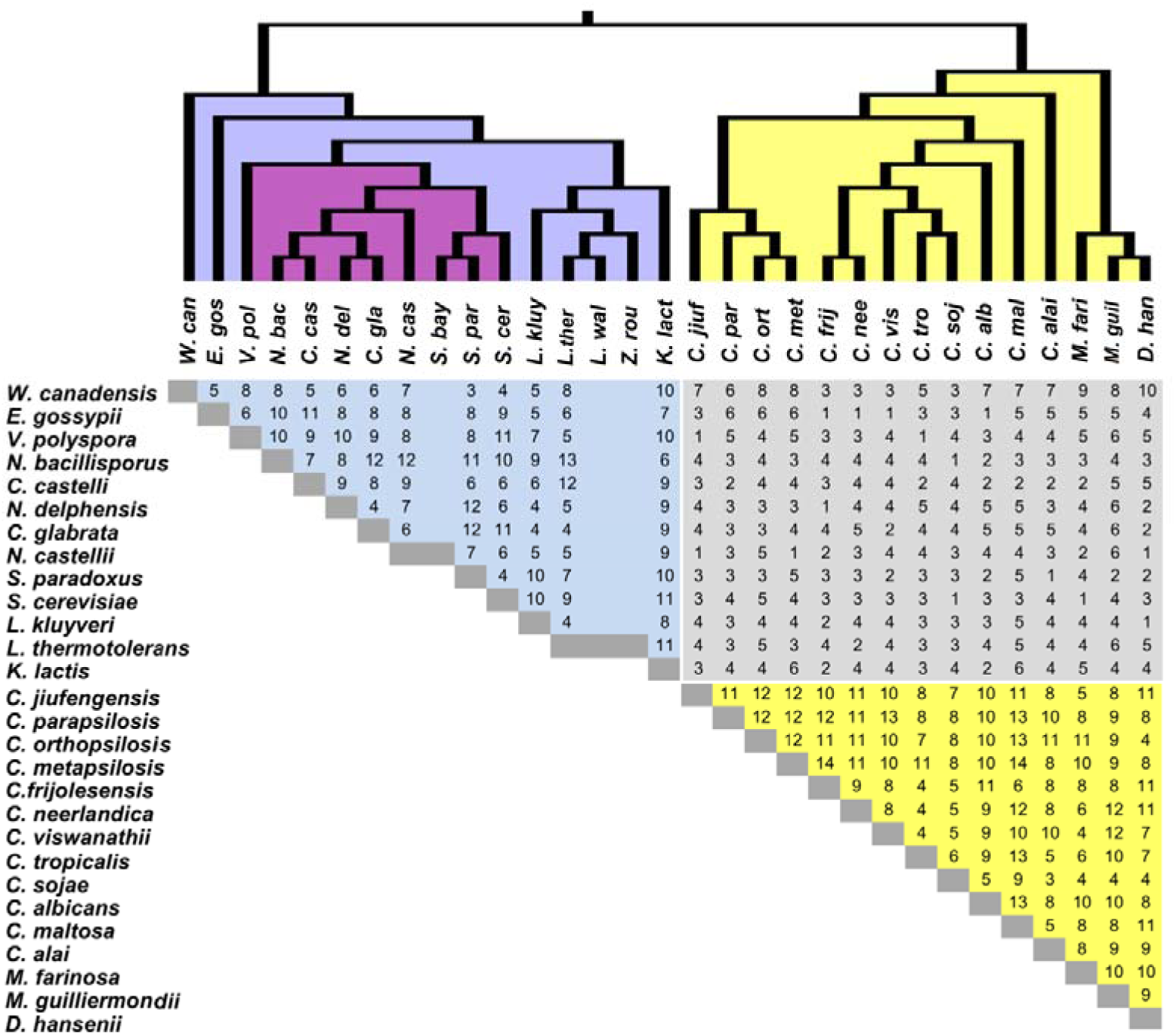
Pairwise similarity of colinear blocks in mitochondrial genomes aligned by Mauve progressive algorithm. Regions of local colinearitycolinear blocks were counted as one conserved block unit. Colored areas (yellow and blue) indicate comparisons within the same clade whereas gray area indicate comparisons between clades. Means ofsyntenic block numbers of each clade are higher than the mean number of conserved blocks between different clades. The nonparametric Mann-Whitney test was used to estimate the significance levels between comparisons.

**Figure 4.**
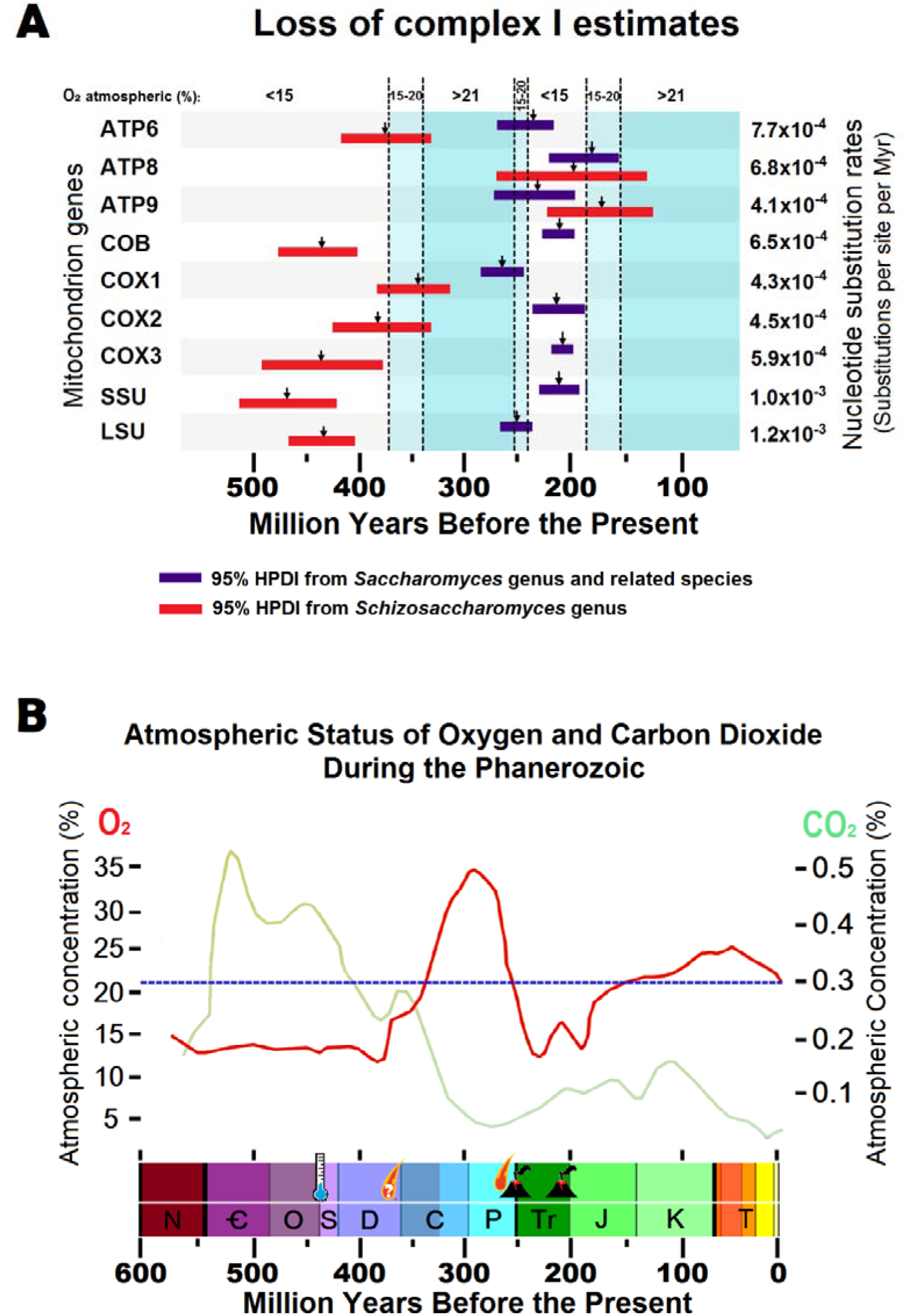
In (A) loss of mitochondrial complex I during the Phanerozoic as estimated by lognormal relaxed molecular clocks and (B) oxygen and carbon dioxide levels and mass extinction events from the Phanerozoic. In (A), red bars indicate 95% HPDI from estimate of the loss of mitochondrial complex I in *Schizosaccharomyces* lineage and blue bars in *Saccharomyces* lineage. Vertical divisions in blue represent oxygenic atmospheric level. In (B) data were compiled from (Berner, 1999; Glasspool and Scott, 2010b; Huey and Ward, 2005; Ward, 2006). The O2 levels are indicated by red lines and CO2 levels by green lines. The five major events extinction events are depicted by symbols representing their putative triggers. The geological periods depicted at the bottom are: Cambrian (Ꞓ), Ordovician (O), Silurian (S), Devonian (D), Carboniferous (C), Permian (P), Triassic (Tr), Jurassic (J), Cretaceous (K) and Tertiary (T).

According to estimative ages based on *ATP8* (265.832 - 136.831 Myr) and ATP9 (265.832 - 136.831 Myr) genes, mitochondrial complex I was lost by *Schizosaccharomyces* more recently,in a period that comprehends the end of Perminian until Jurassic, the last great hypoxic period (Table 4 and Figure 4). This period estimated includes two mass extinction event:Permian–Triassic extinction (252 Myr) and End Triassic mass extinction (199 – 214 Myr). The Permian–Triassic mass extinction episode, occurred around 252 Myr, was considered the worst of the five mass extinctions, due 95% of all species, including marine and terrestrial, were eradicated (Benton, 2015; Huey and Ward, 2005; Ward, 2006). The event was, possibly, caused by a bolide impact that initiated a volcanic flood emanating from the Siberian Traps (Benton, 2015; Kaiho et al., 2001; Wake and Vredenburg, 2008). Consequently, there was a severe decline of oxygen (Huey and Ward, 2005). The hypoxic environment persisted along million years until the next mass extinction event, in the End Triassic. During Triassic-Jurassic (Tr-J) boundary (Figure 4B), around 200 Myr ago, occurred the Central Atlantic Magmatic Province (CAMP) volcanic eruptions, one of the strongest volcanic episodes in Earth’s history (Hesselbo et al., 2002; Pálfy et al., 2000). Emission of voluminous volcanic gases lead to increased CO_2_ and decreased O_2_ and coincided with climatic crisis and abrupt decline of marine and terrestrial flora and fauna (Hesselbo et al., 2002), leading to the loss of the whole class of conodonts (Stanley, 2009), and the extincion of 48% invertebrate species (Keller, 2005).

While most of our estimates indicate that *Schizosaccharomyces* may have lost the complex I in the first half of Phanerozoic, the loss of mitochondrial complex I by *Saccharomyces* was dated exclusively in second half, between Middle Permian until Middle Jurassic periods (Table 4 and Figure 4). As well as discussed above, this hypoxic period coincides with End Permian mass extinction and End Triassic mass extinction events. The persistent oxygen low levels associated with events that provide abrupt changes in climate and environment may have favored the loss of respiratory complex I. During hypoxia, our results allow us to supposed that the complex I genes were loss as an economical molecular strategy. Declined oxygen levels may have been consumed even when mitochondrial respiratory chain from yeasts became defective. Without complex I to transfer electrons using NAD cofactor, succinate dehydrogenase, the mitochondrial complex II, transfers electrons reducing FAD cofactor, promoting the execution of respiratory chain (Rutter et al., 2010). However, decreased oxygen levels induces production of ROS (Pitkanen and Robinson, 1996) and, consequently, contribute to damages and instability on mitochondrial and nuclear genomes (Kaniak-Golik and Skoneczna, 2015; Rasmussen et al., 2003; Stuart et al., 2006).

The absence of complex I could have affected yeasts mitochondrial-nuclear genomic stability. Could WGD event have occurred as result of nuclear instability? Two estimates for WGD, based on *ATP9* (246.297 - 166.663 Myr) and *COX2* (205.566 - 121.007 Myr) genes, overlapped with the estimates of the loss of complex I by *Saccharomyces*, *ATP9* (266.790 - 197.769 Myr) and *COX2* (239.12 - 190.512 Myr). These results permit us to presume the WGD as a consequence of mitochondrial instability caused by the loss of complex I. On the other hand, the most of WGD estimates are not overlapped with loss of complex I event, although some estimate pointed both events as subsequent episodes (*ATP6* and *COX3*). Our WGD estimative based on nuclear 18S rRNA (134.899 - 108.413 Myr) is compatible with the WGD estimate of 150 to 100 Myr (Wolfe and Shields, 1997). *ATP8*, *COB*, *COX2* and SSU mitochondrial genes also comprehends part of the classical estimative from Wolfe and Shields. Therefore, based on molecular clock estimate, WGD occurred after the loss of complex I by *Saccharomyces*.

Hyperoxic or normoxic conditions might have alleviated the negative selection on respiratory systems of yeasts, which by this time had a fermentative metabolism well developed since the availability of land plant fruits (Dashko et al., 2014). *S. cerevisiae* obtain energy preferentially by fermentation, and they do not exhibit the Pasteur effect, where fermentation is inhibited by oxygen. Therefore, complex I defective mutants in the *S. cerevisiae* lineage under hypoxia, where fermentation is preferred, would have lost the complex I. With the subsequent increase on oxygen levels, the adaptations for respiration might have pressured these organisms to use the nuclear *NDE1* and *NDI1* genes to replace the absent to classical complex I to start the respiratory chain (Luttik et al., 1998; Postmus et al., 2011b). Despite the evolutive distance between complex I negative species *Schizosaccharomyces pombe* and *S. cerevisiae* (Melo et al., 2004), the *NDE1* and *NDI1* genes have a pattern of convergent evolution, as observed by the comparison of *NDE1* and *NDI1* genealogies (Figures 5A and 5C). Differently than 18S rRNA phylogeny (Figures 5B and 5D), *Sz. pombe* and *S. cerevisiae* were placed closest on phylogenetic trees based on *NDE1* and NDI1. *NDE1* and *NDI1* genes are not related to the original mitochondrion encoded NADH genes, and are present in complex I positive and complex I negative lineages. This can be explained because in complex I negative lineages (*Schizosaccharomyces* and *Saccharomyces*) the *NDI* and *NDE1* genes have to substitute the function of missing mitochondrial encoded complex I genes. This functional complementation is however not necessary in complex I positive lineages. Other Saccharomycetales subgroups that don’t have strong fermentative metabolism might have devised other strategies to cope with hypoxia that could not allow for the loss of the complex I, as is the case of *Candida albicans* (Rozp□dowska et al., 2011). These might include increased respiratory rates and uncoupling (Cavalheiro et al., 2004). In the *S. cerevisiae* pre-WGD clade it is important to notice that *Kluyweromyces lactis* does not have the Crabtree effect and cannot use ethanol (Liti et al., 2001). NADH gene remnants were not observed in complex I negative mitochondrial DNA (data not shown) which is consistent with a scenario of ancient loss of complex I, which occurred most likely by deletions of inactive genes.

**Figure 5.**
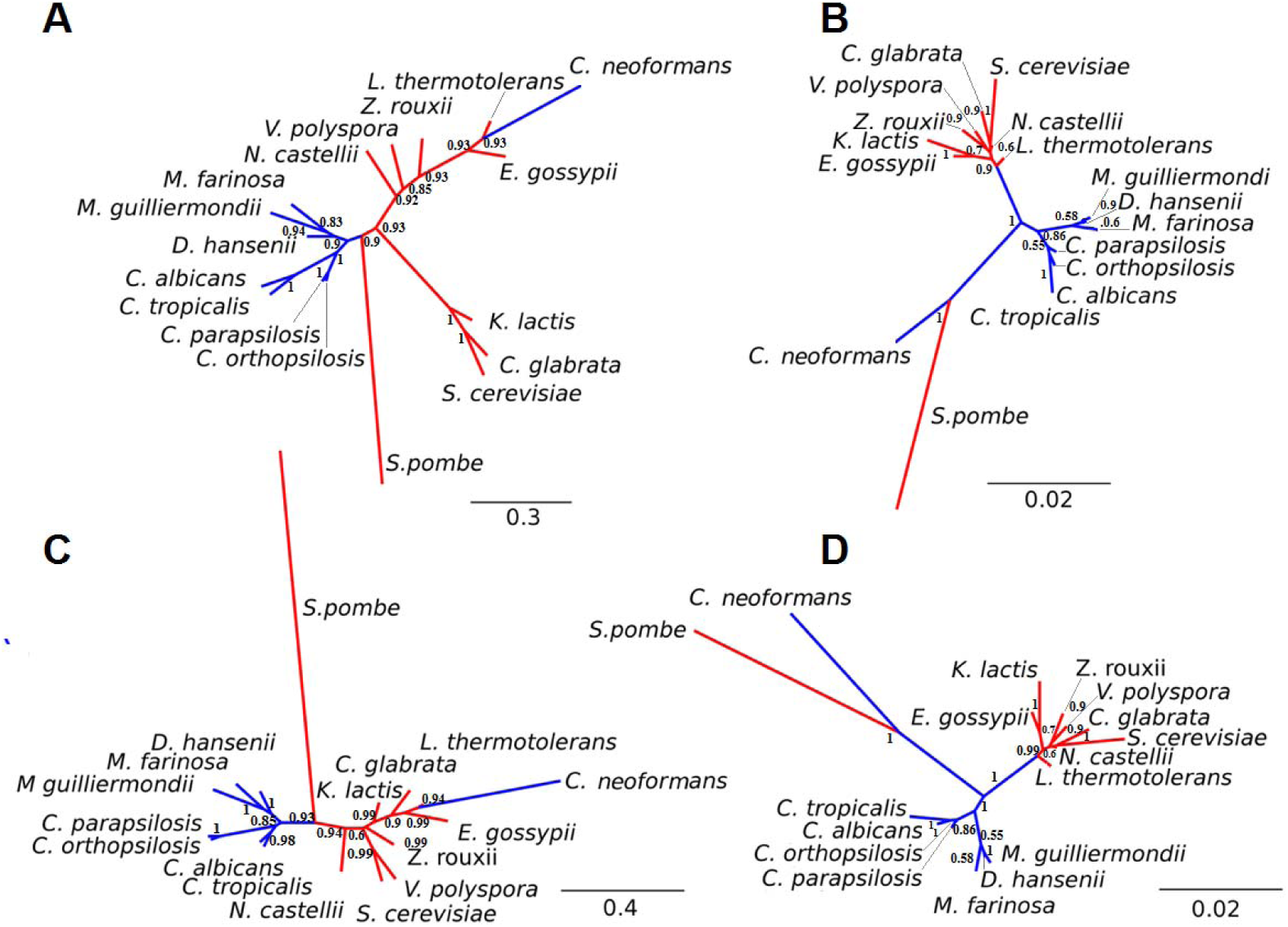
Gene tree of NDI (A) and corresponding 18S rRNA 1 phylogeny (B) and gene tree of *NDE1* (C) and the corresponding 18S rRNA phylogeny (D) in Saccharomycetales. In red are the complex I negative taxa and blue the complex I positive taxa. In both NDI and *NDE1* phylogenies the *Schizosaccharomyces* branch is more closely related to other complex I negative clades as compared to the 18S rRNA trees where *Schizosaccharomyces* is distantly related to *S. cerevisiae*. This can be explained because in complex I negative lineages (Schizosaccharomyces and Saccharomyces) the NDI and *NDE1* genes have to substitute the function of missing mitochondrial encoded complex I genes. This functional complementation is however not necessary in complex I positive lineages (e.g. *C. albicans*).

## 4. Conclusion

The mitochondrial phylogenomic, synteny analysis and molecular clock estimates here presented confirm that the loss of complex I occurred before, maybe subsequently, the WGD and independently in *S. cerevisiae* and *Schizosaccharomyces*. Correlations of estimated dates and variations of Phanerozoic atmospheric oxygen concentrations suggest that, the loss of complex I by yeast may have occurred during hypoxic periods, possibly associated with mass extinction events. The *Saccharomyces* lineage might have lost the mitochondrial complex I thanks to hypoxia caused by the End Permian or the End Triassic mass extinction events, while *Schizosaccharomyces* lineage might have lost the mitochondrial complex I associated with Late Ordovician or Late Devonian mass extinction events. Both periods with low oxygen environment where negative selection was probably alleviated on complex I defective mutants under hypoxia where fermentation would have been preferred. The loss of complex I and its pressure for necessary substitution by nuclear gene products during the oxygen increase periods might have, at least in part, contributed to genome instabilities associated with WGD as in the case of *S. cerevisiae* and its relatives. We conclude that supertrees and relaxed molecular clocks suggest losses of yeast mitochondrial complex I in Late Devonian and Triassic-Jurassic boundary.

## Supporting information

