## Supplementary figures and images for "Evolution of pathogenic and nonpathogenic yeasts mitochondrial genomes inferred by supertrees and supermatrices with divergence estimates based on relaxed molecular clocks"

### Sup_figure1_15S.tiff

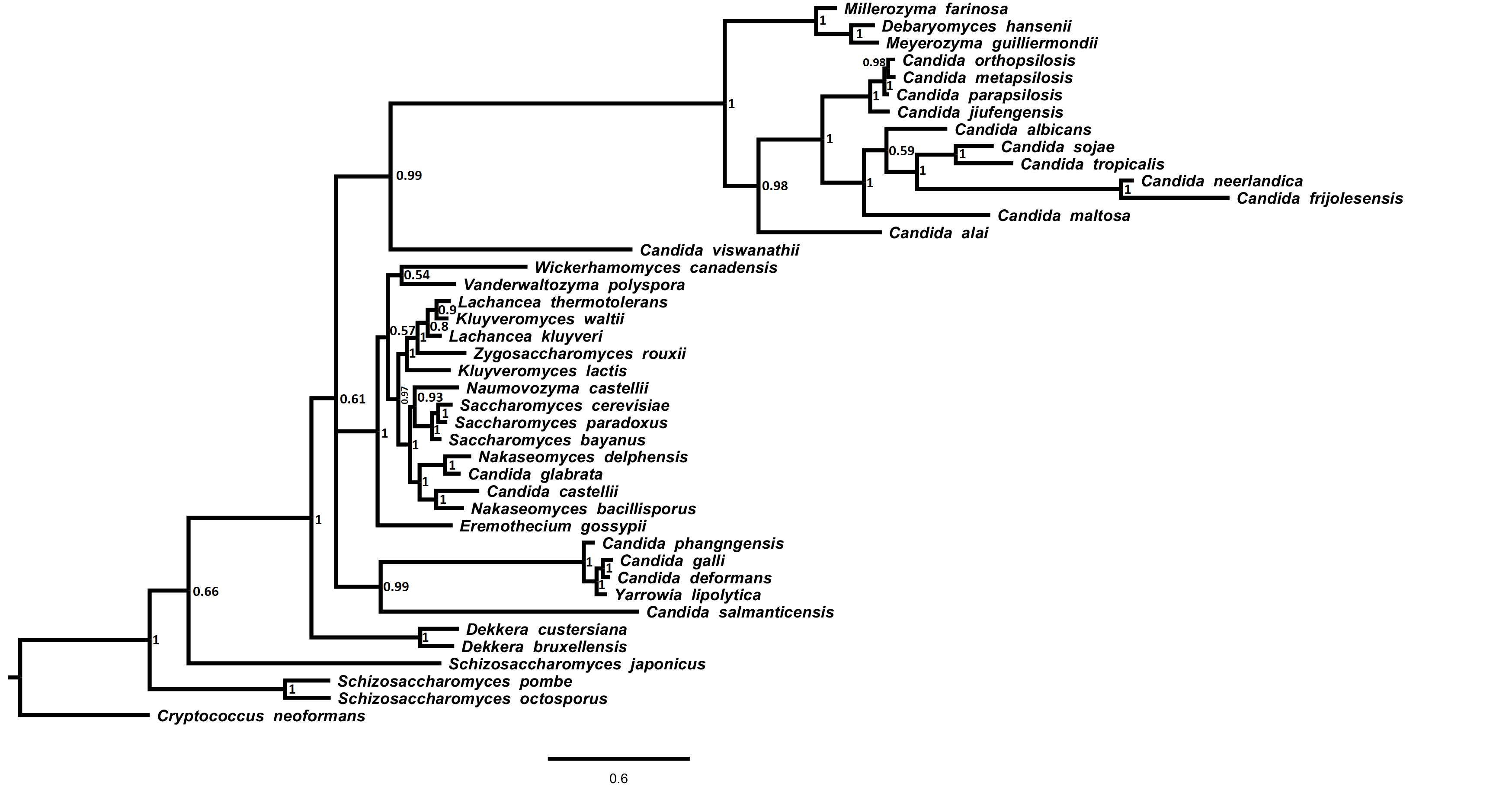

### Sup_figure2_21S.tiff

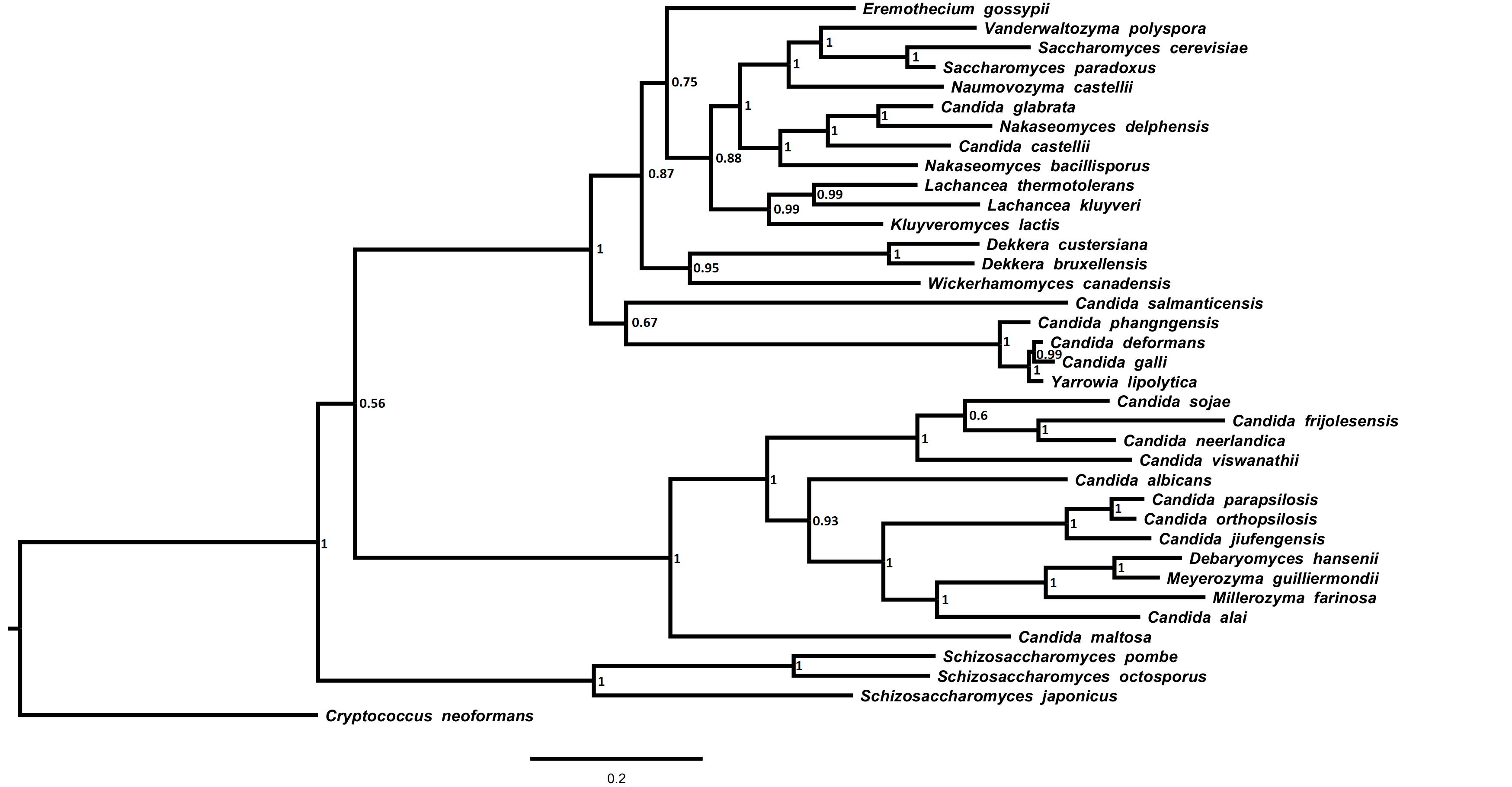

### Sup_figure3_ATP6.tiff

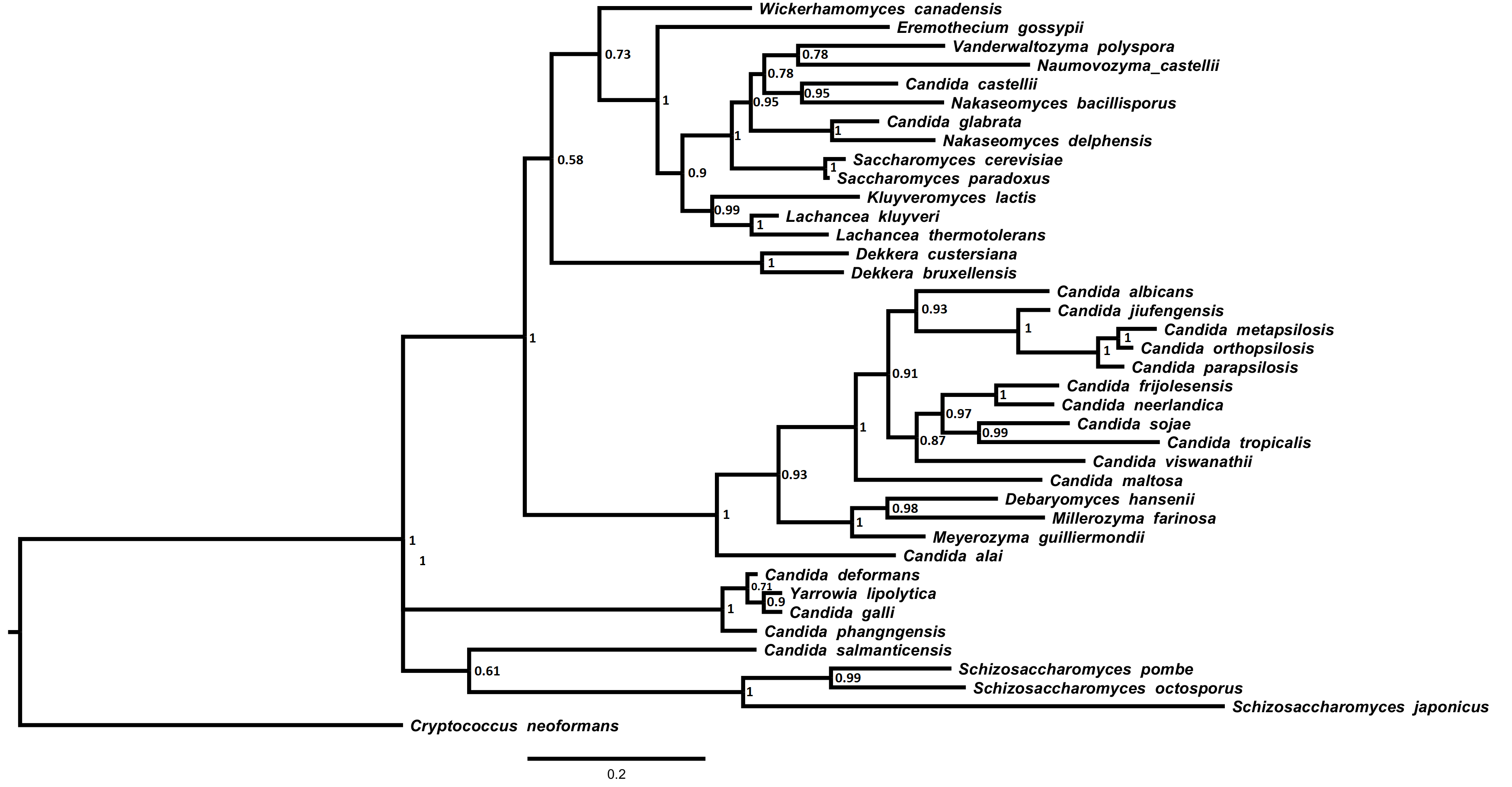

### Sup_figure4_ATP8.tiff

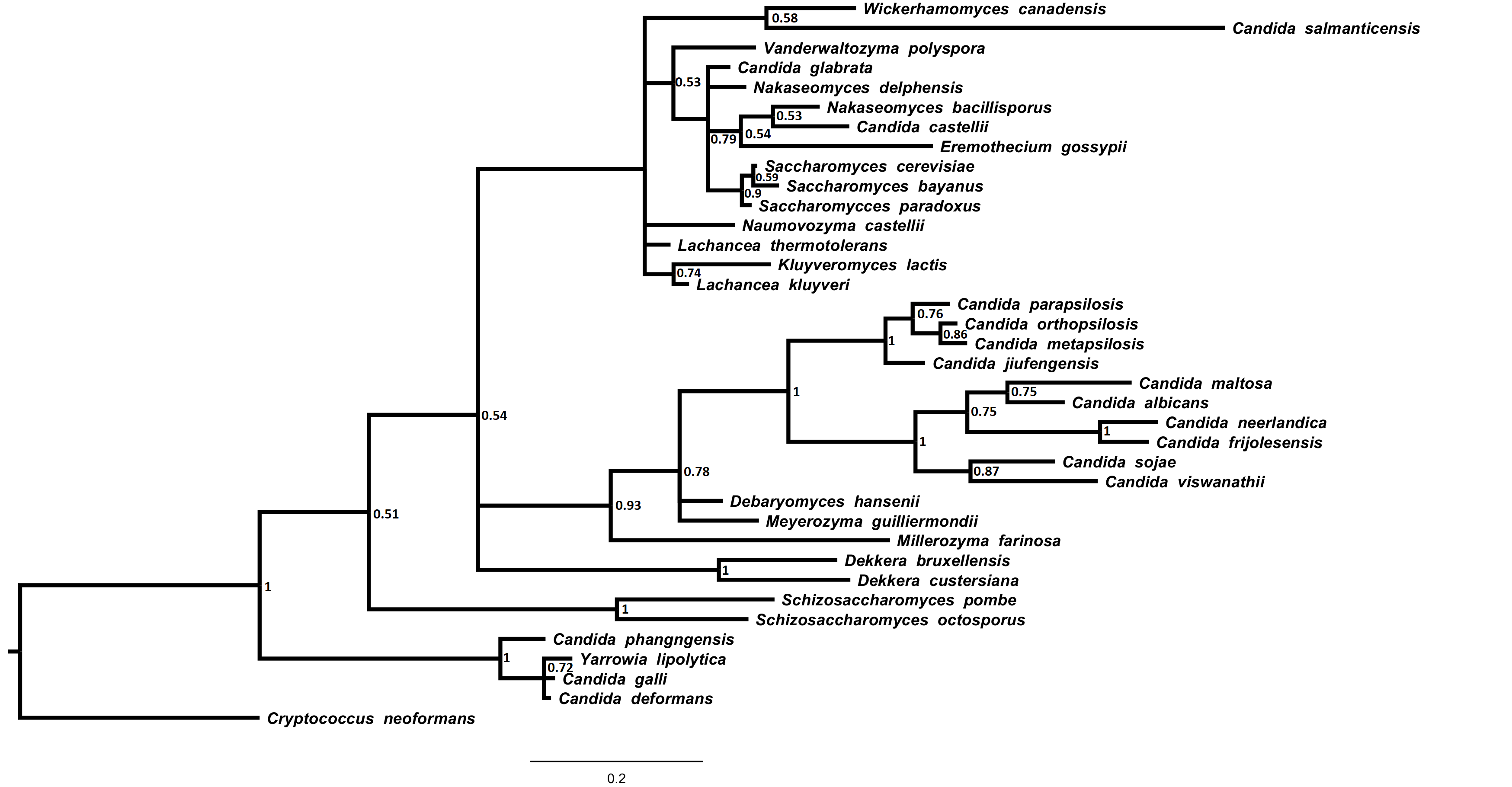

### Sup_figure5_ATP9.tiff

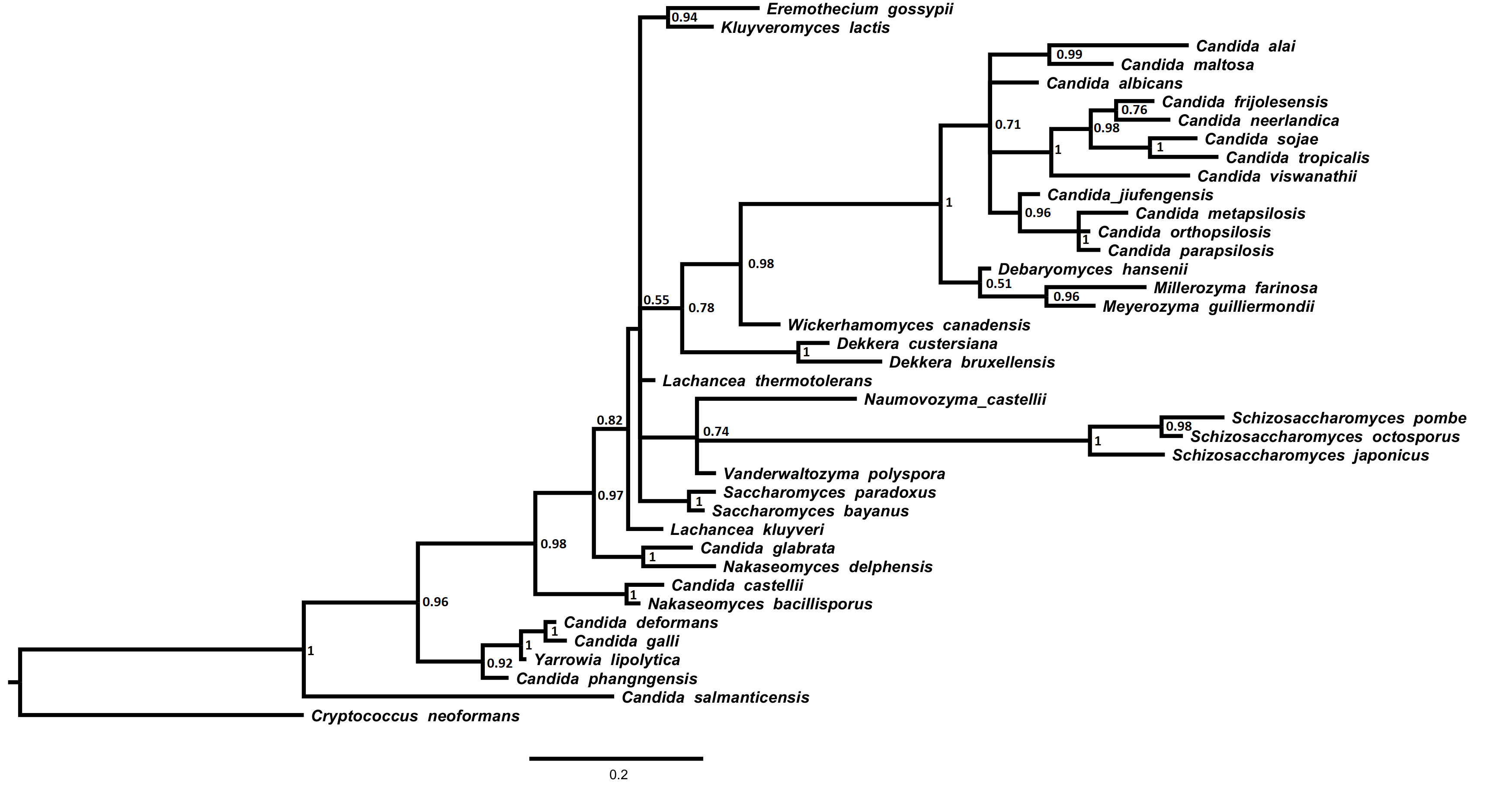

### Sup_figure6_COB.tiff

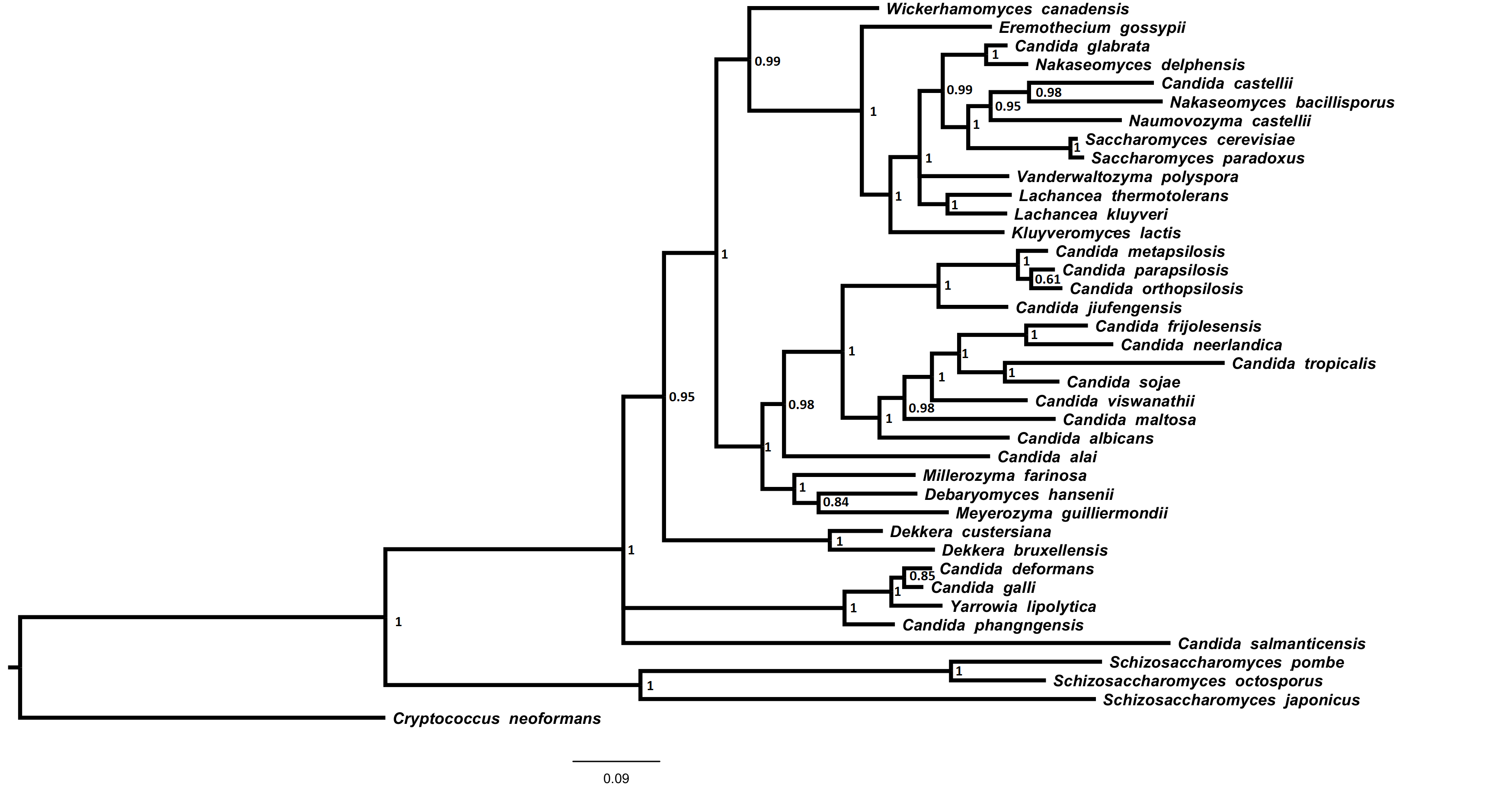

### Sup_figure7_COX1.tiff

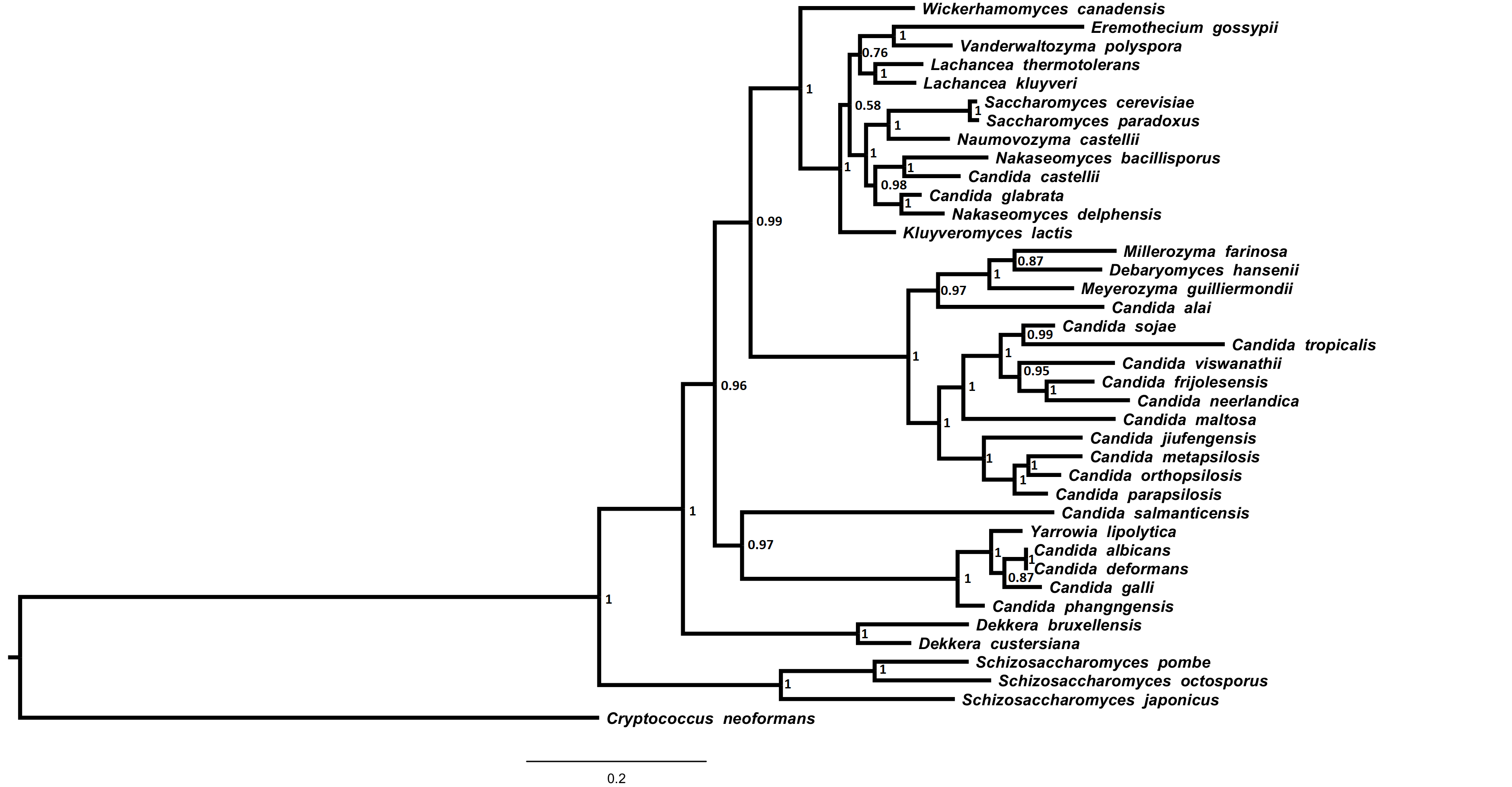

### Sup_figure8_COX2.tiff

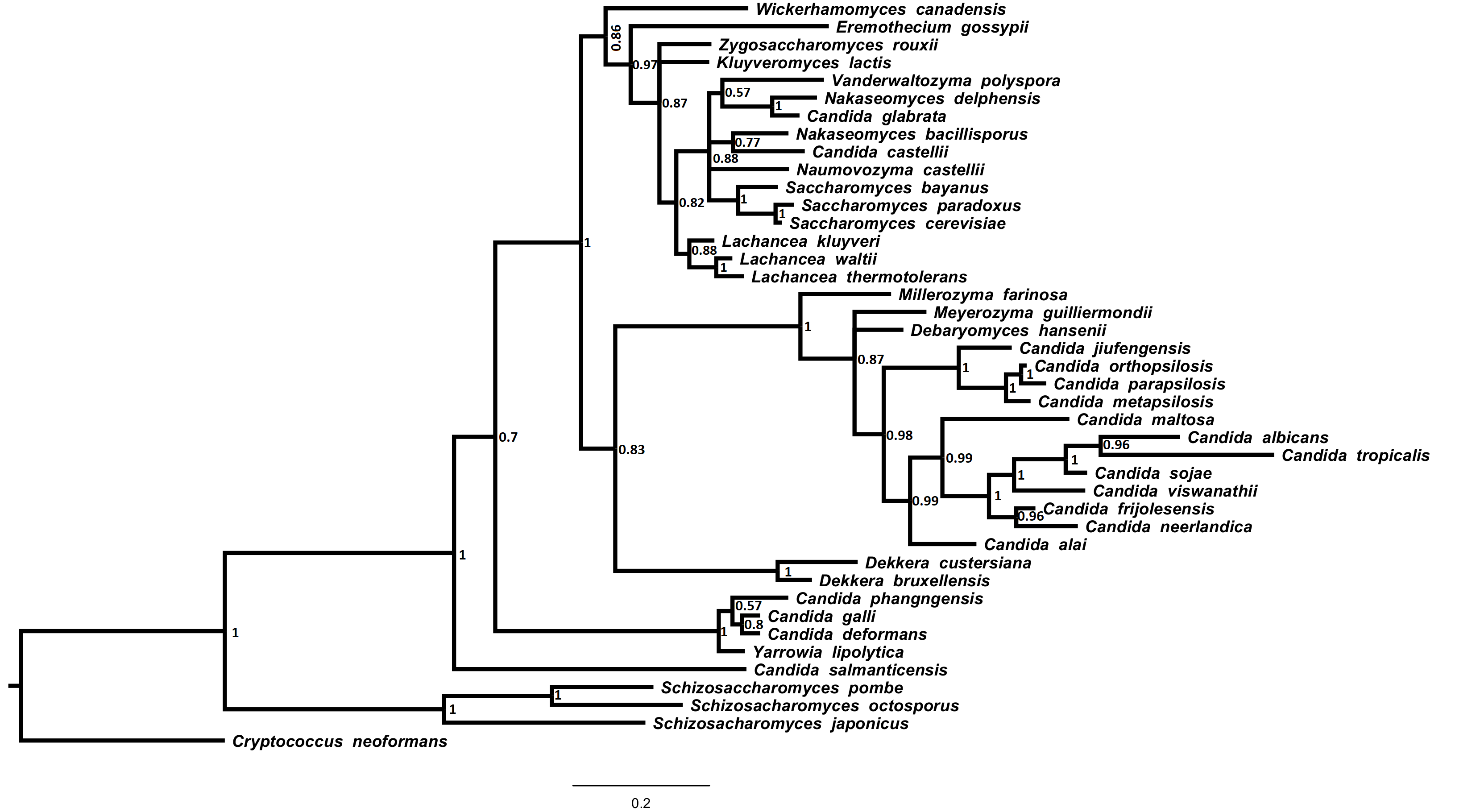

### Sup_figure9_COX3.tiff

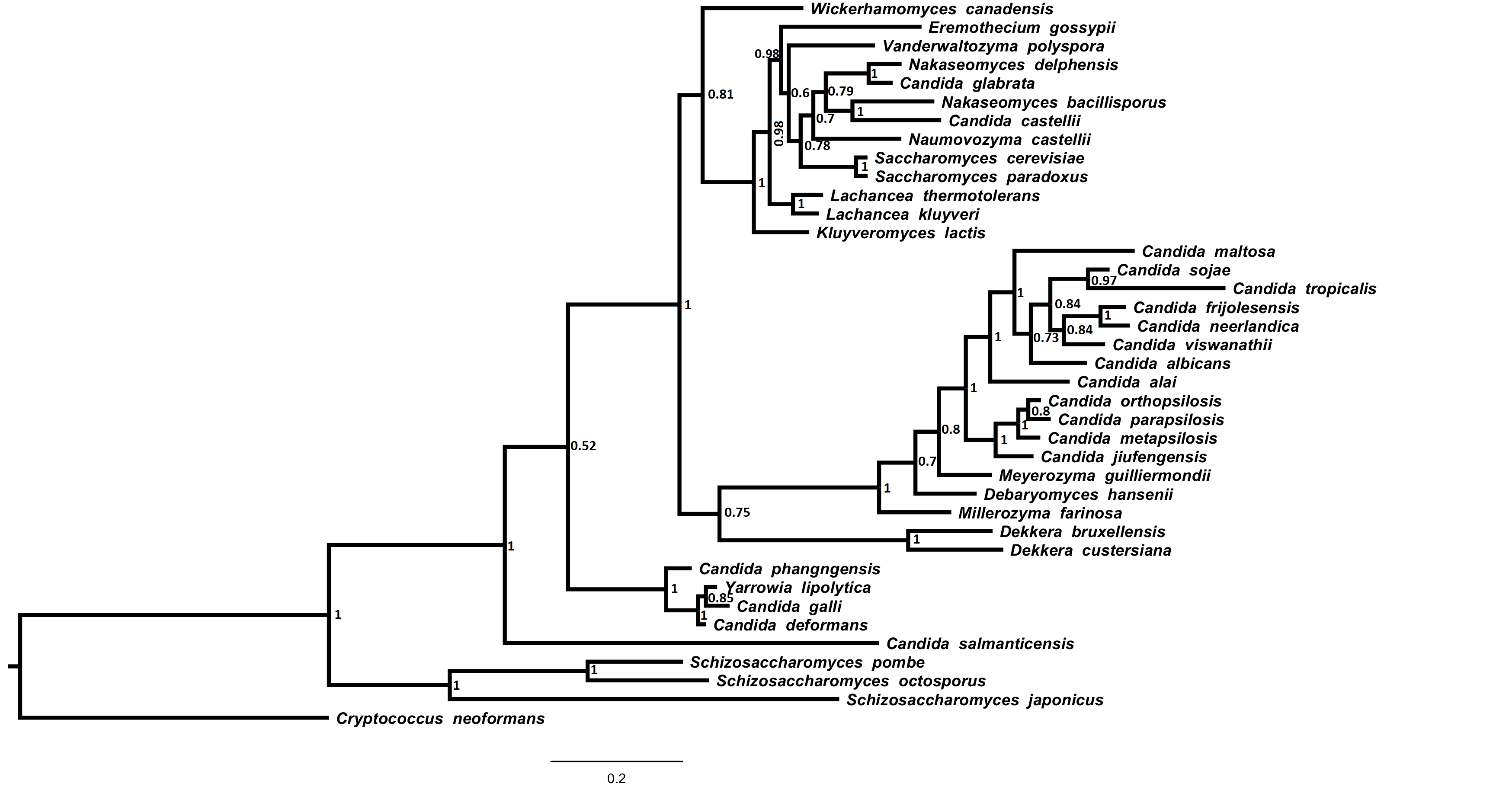

### Sup_figure10_NAD1.tiff

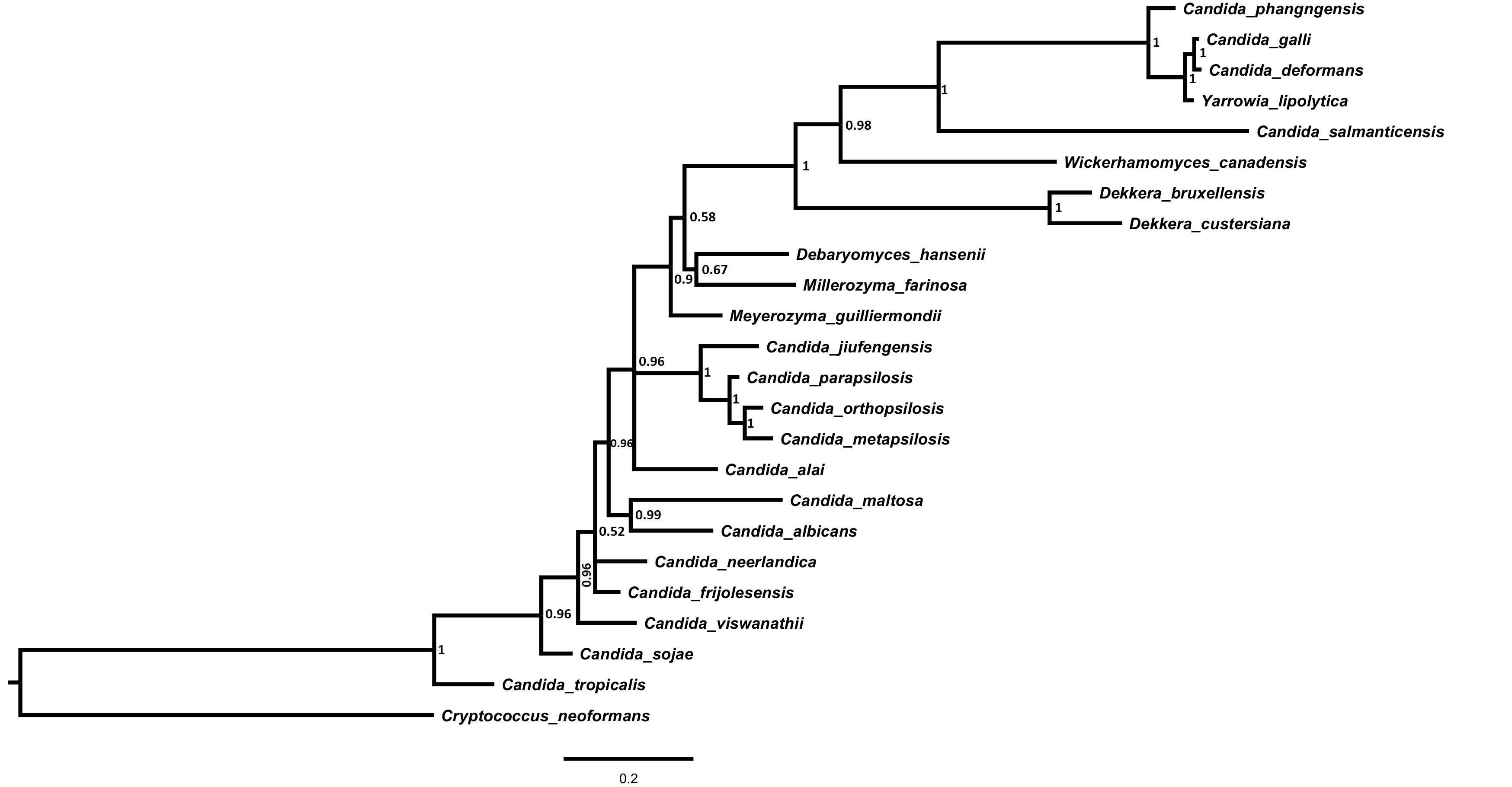

### Sup_figure11 -NAD2.tiff

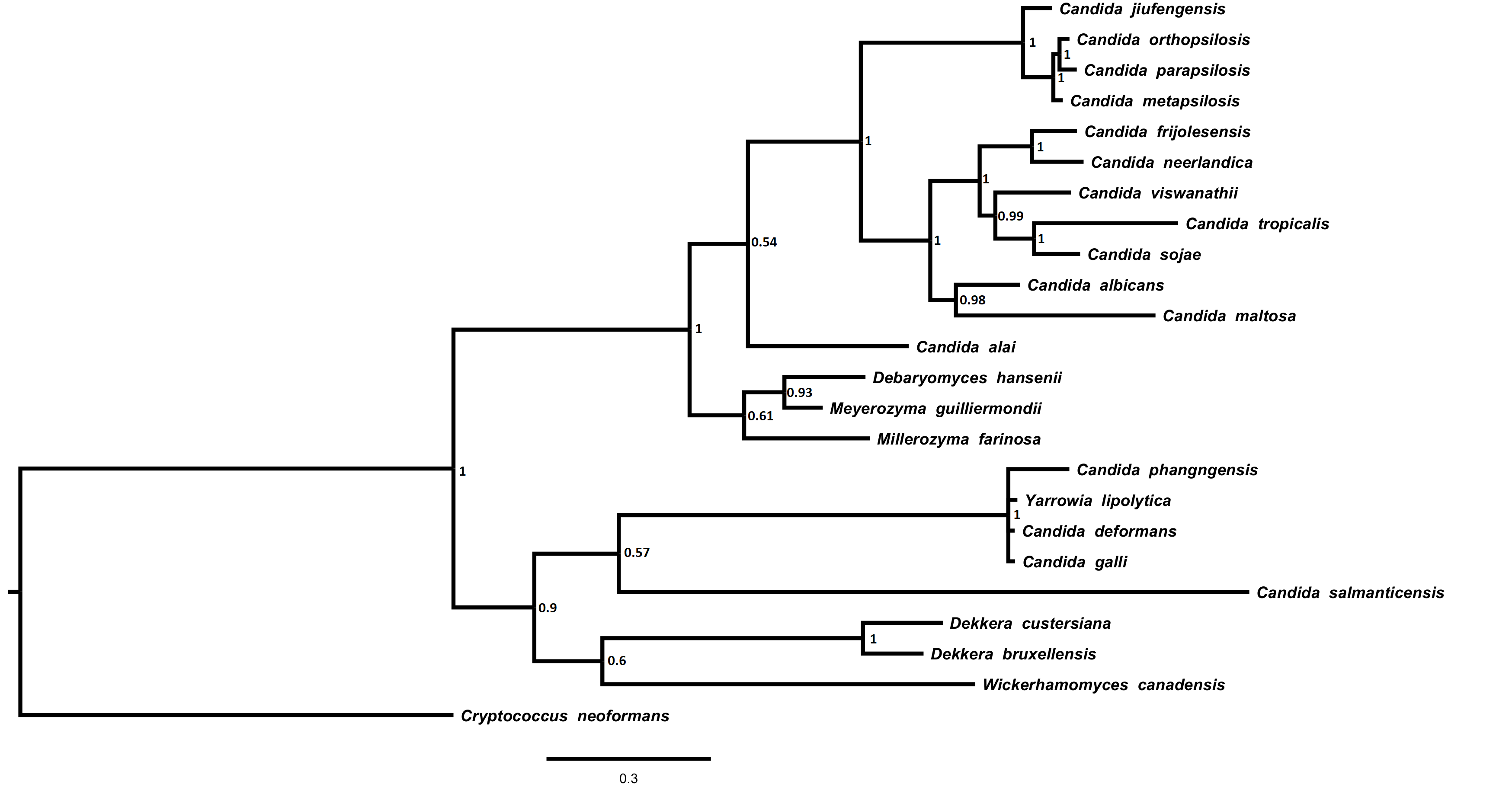

### Sup_figure12_NAD3.tiff

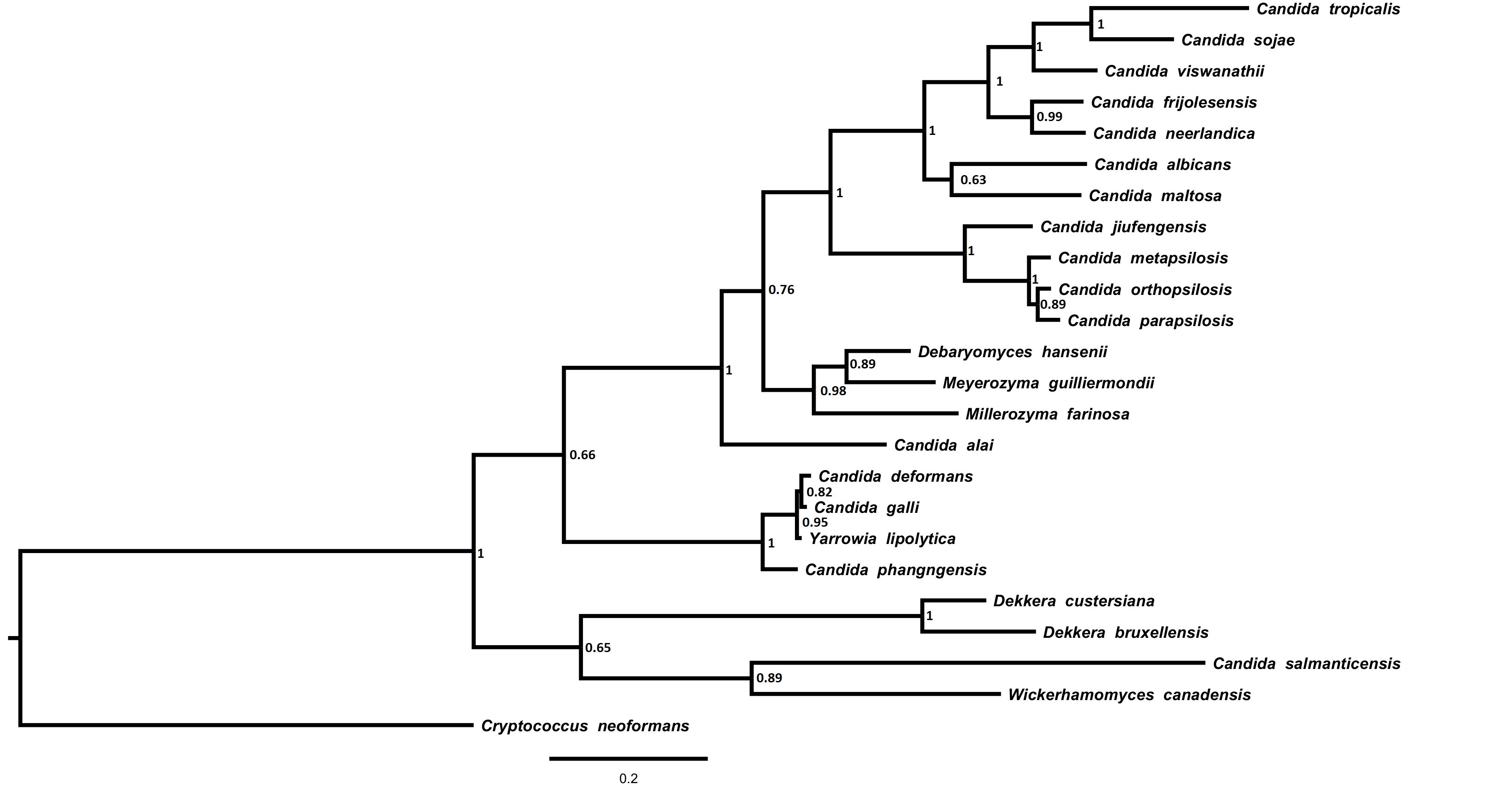

### Sup_figure13_NAD4.tiff

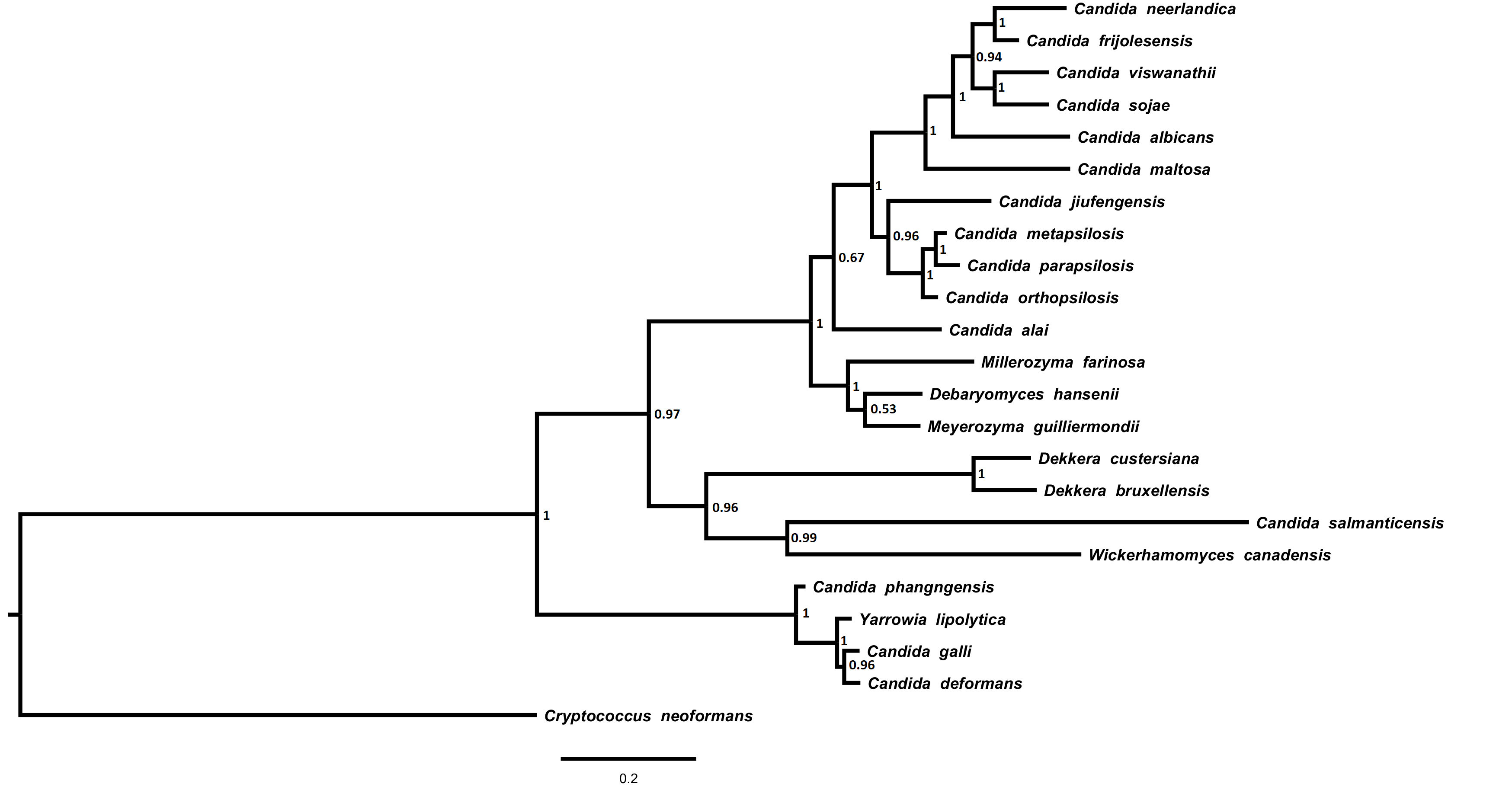

### Sup_figure14_NAD4L.tiff

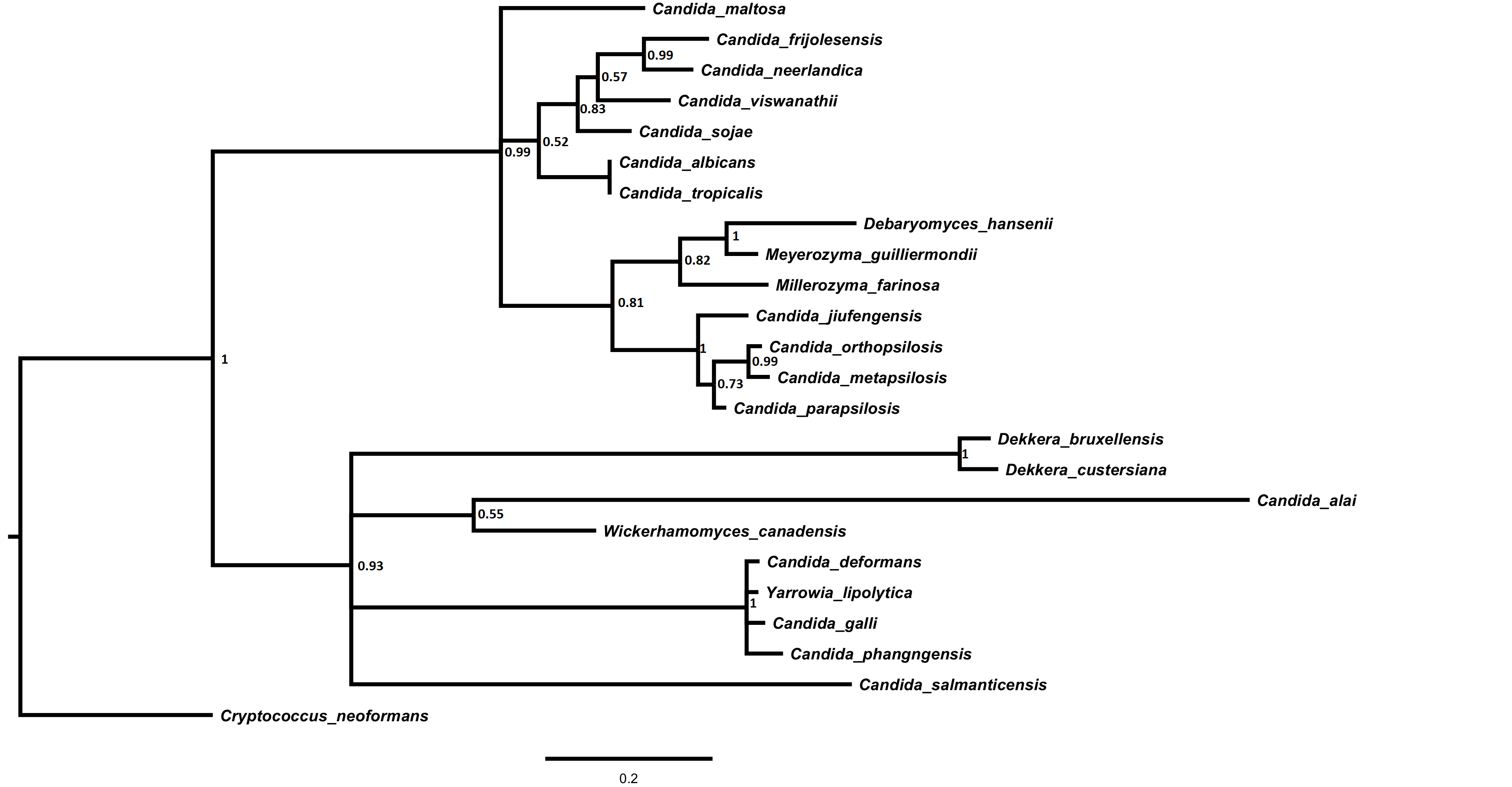

### Sup_figure15_NAD5.tiff

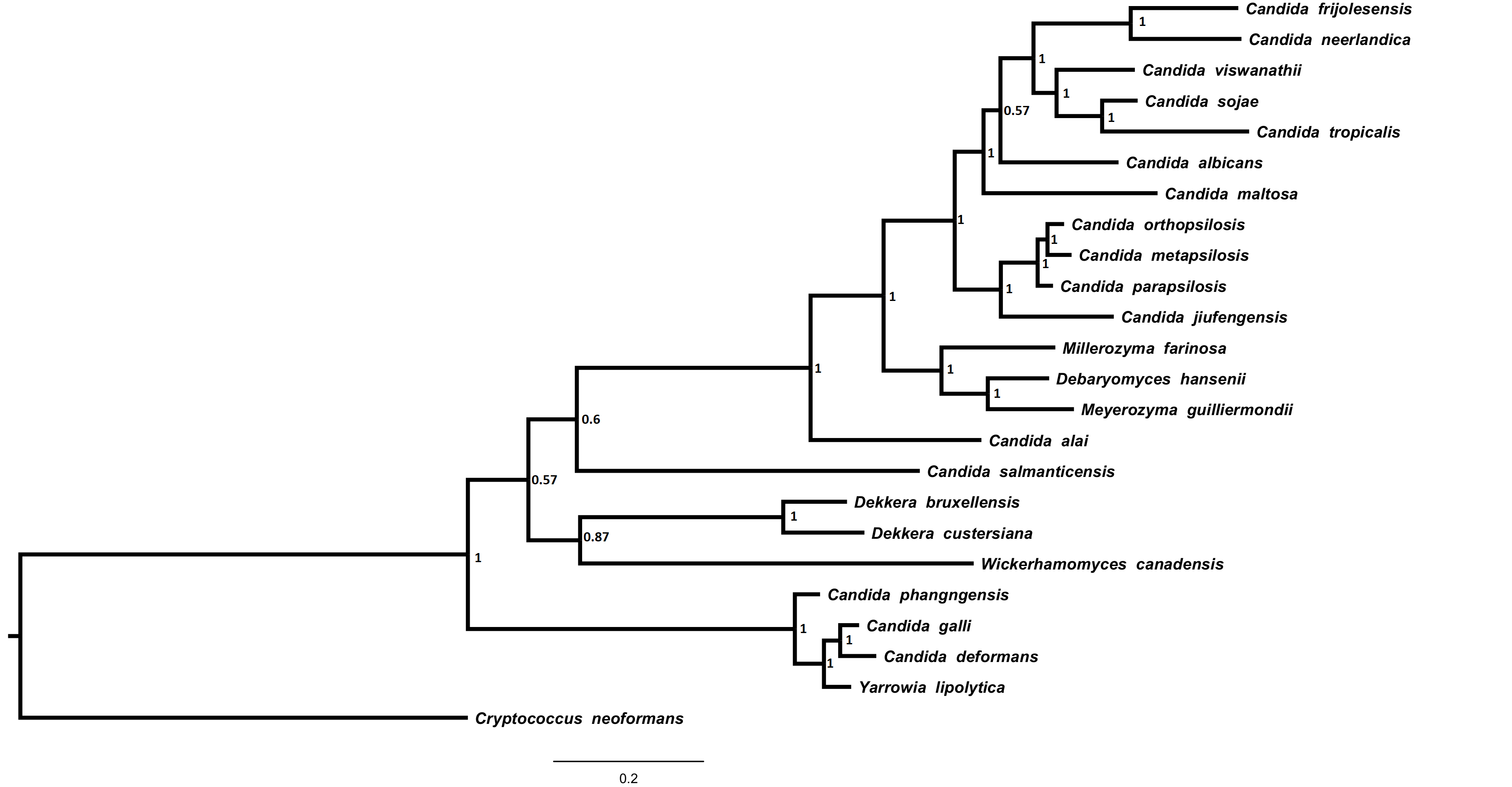

### Sup_figure16_NAD6.tiff

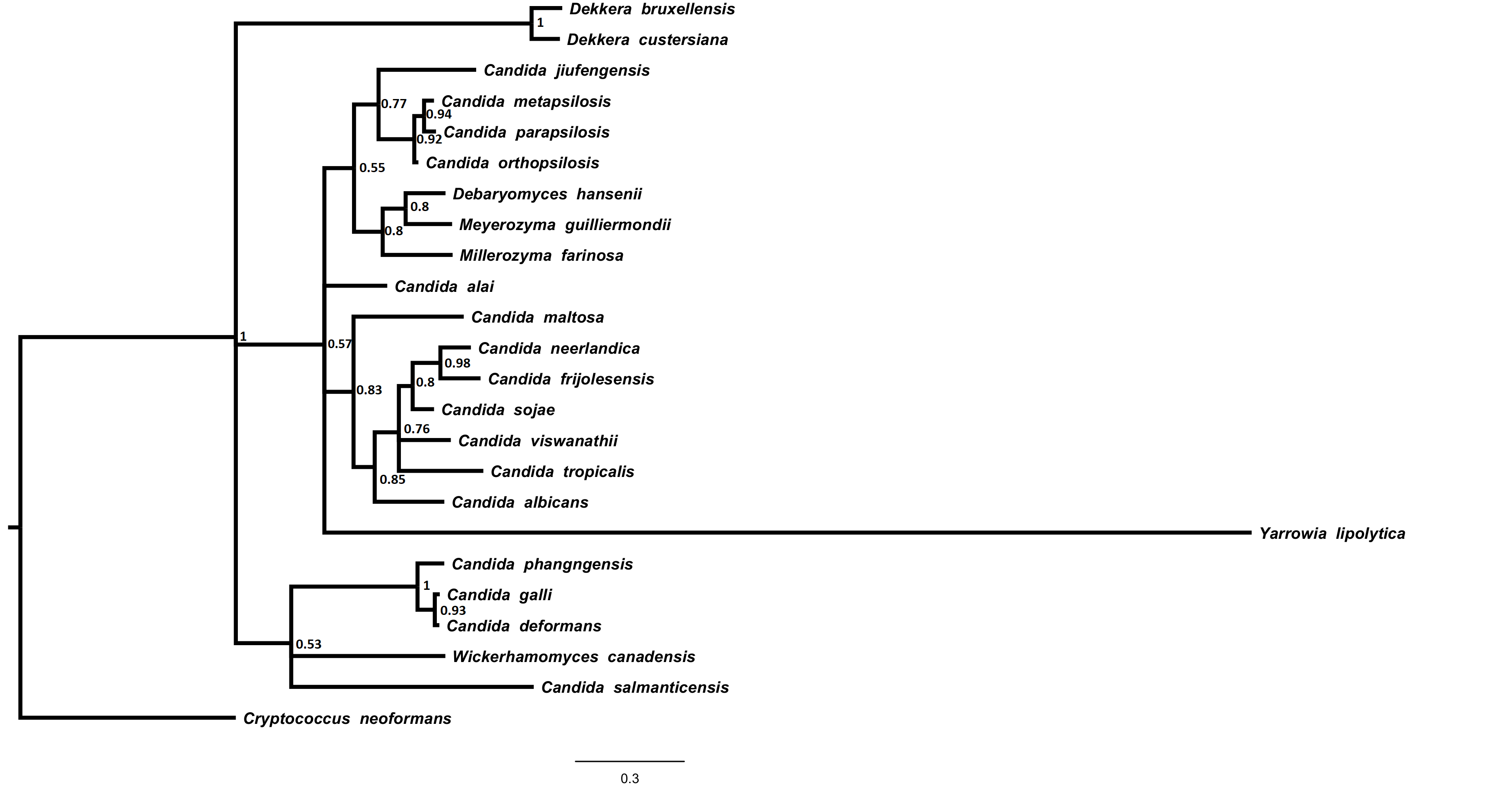

### Sup_figure17_RFSupertree.tif

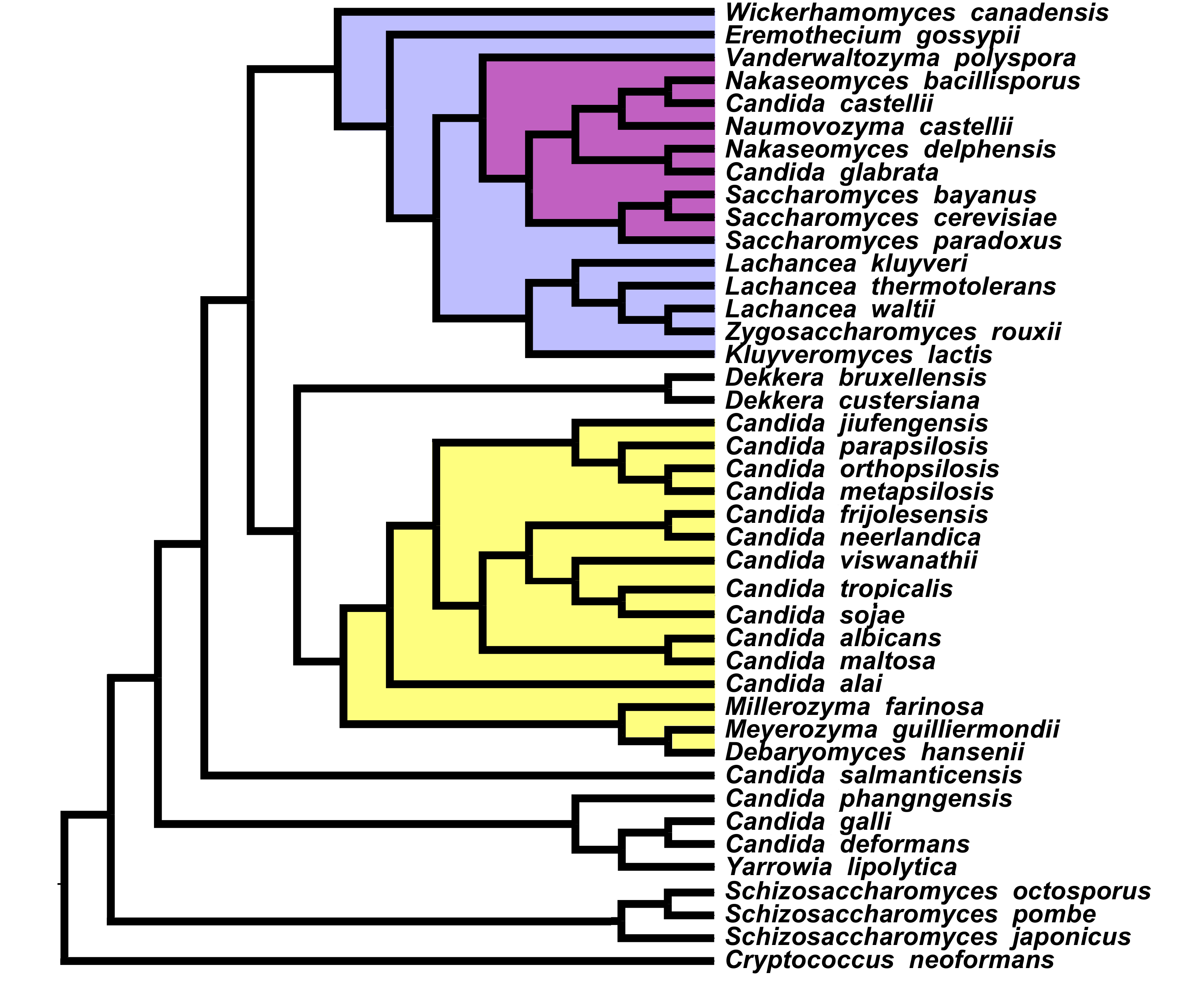

### Sup_figure18_MRPSupertree.tif

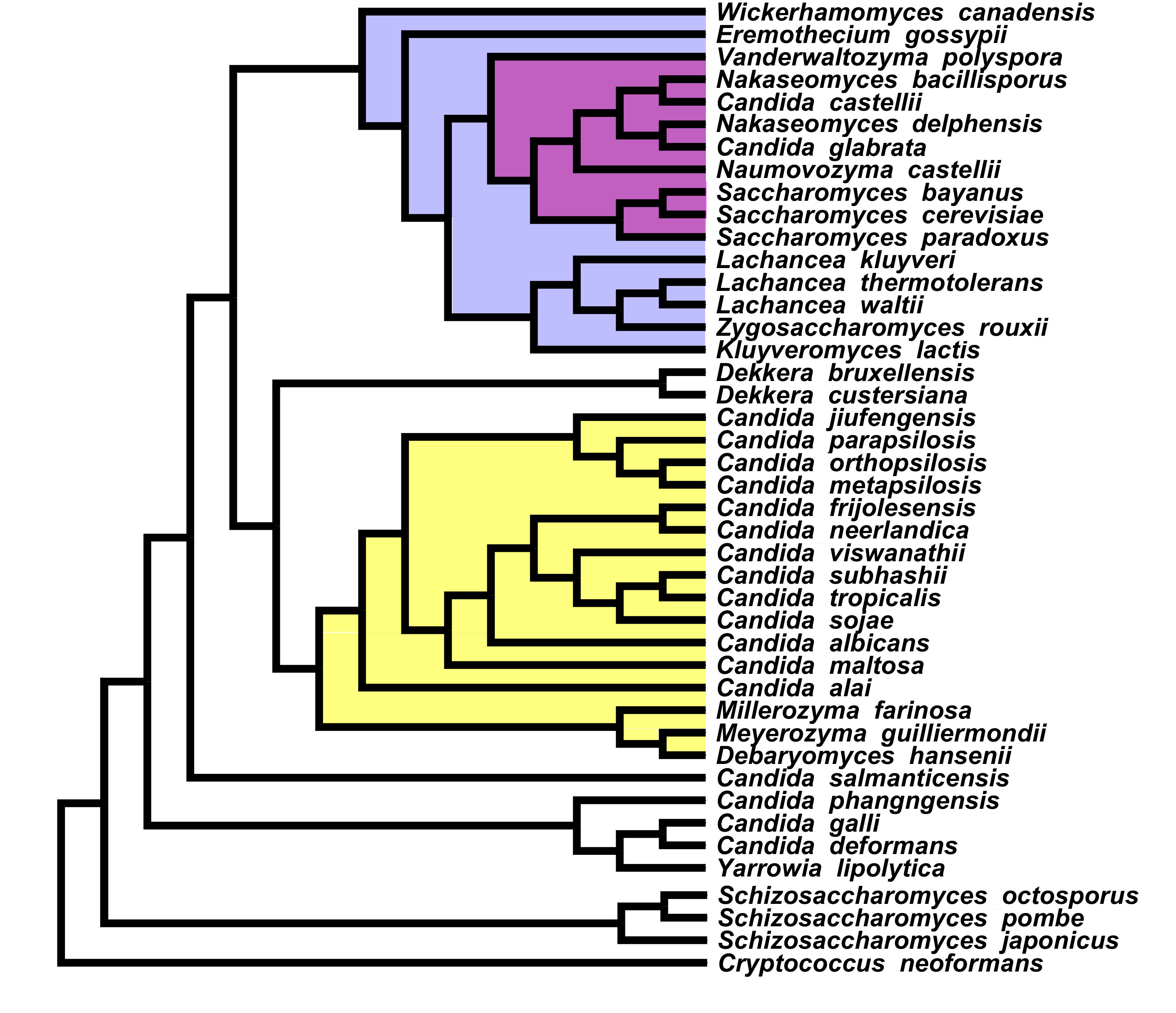

### Sup_figure19_Supermatrix.tif

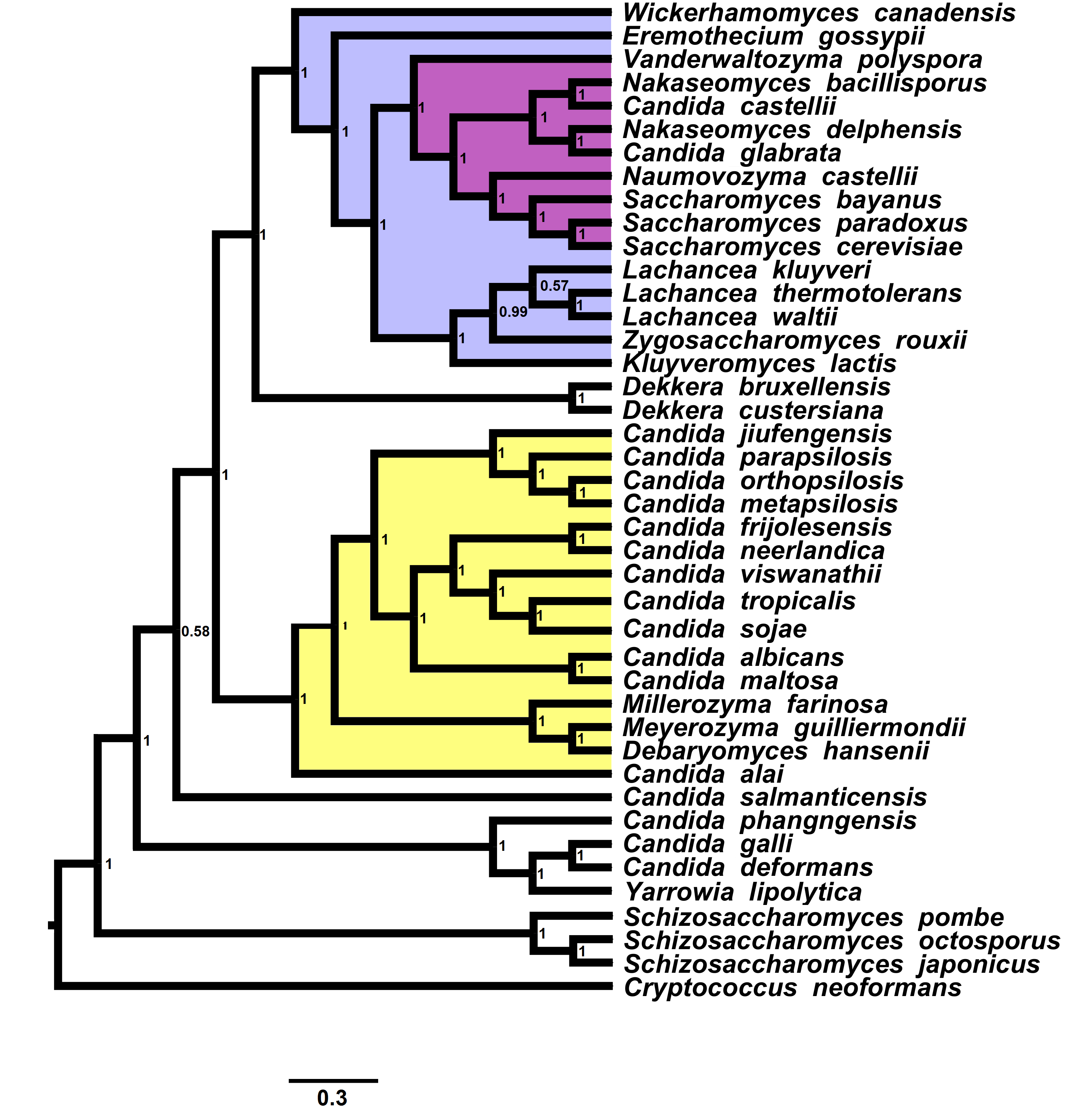

### Sup_figure20_18S.tiff

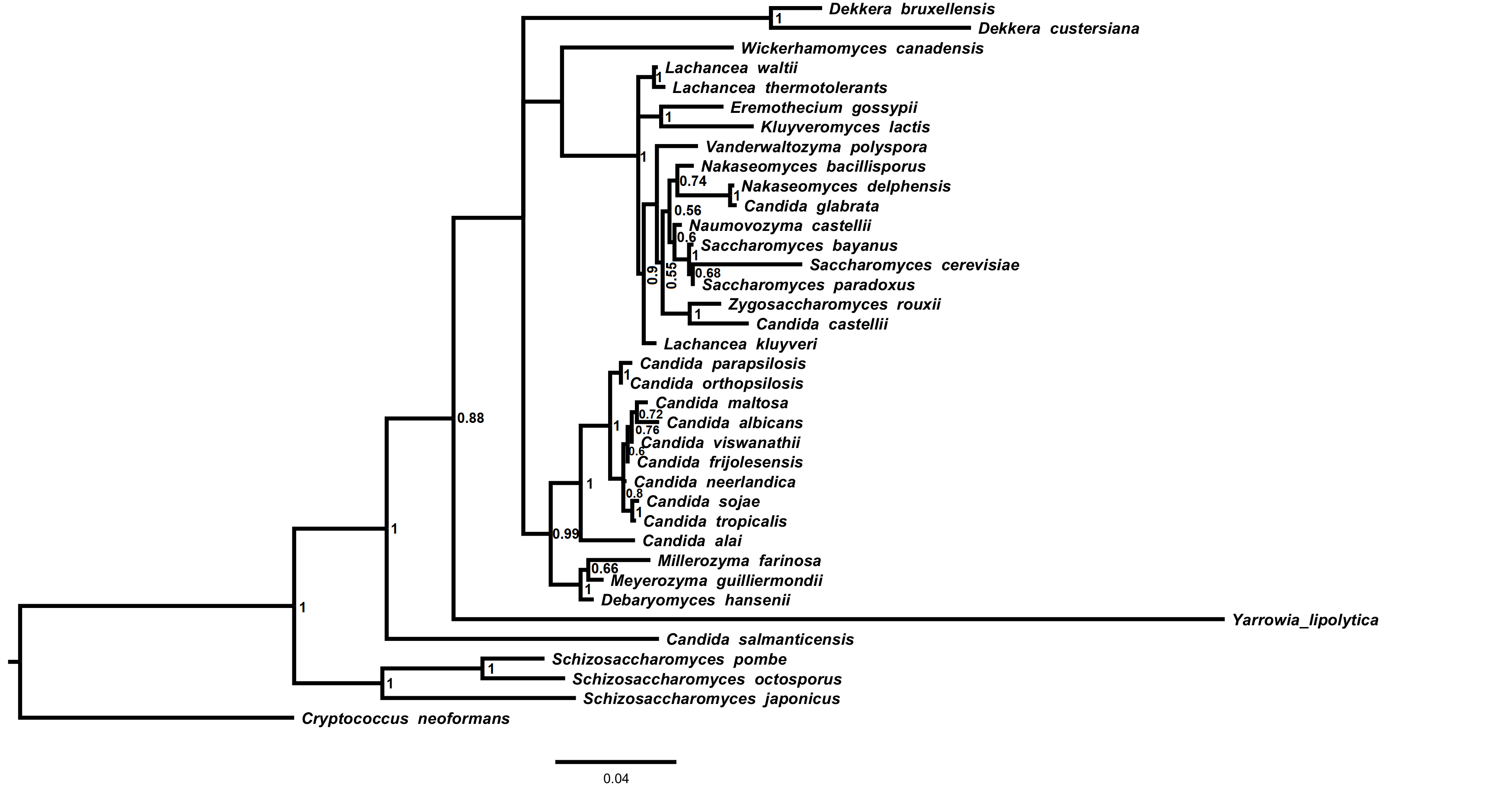
